# An Essential Adaptor for Apicoplast Fission and Inheritance in Malaria Parasites

**DOI:** 10.1101/2025.04.12.648511

**Authors:** James Blauwkamp, Krithika Rajaram, Sophia R. Staggers, Oliver Harrigan, Emma H. Doud, Wei Xu, Hangjun Ke, Sean T. Prigge, Stella Y. Sun, Sabrina Absalon

## Abstract

Blood-stage *Plasmodium falciparum* parasites rely on a non-photosynthetic plastid, the apicoplast, for survival, making it an attractive target for antimalarial intervention. Like the mitochondrion, the apicoplast cannot be generated *de novo* and must be inherited by daughter parasites during cell division. This inheritance relies on coordinated apicoplast positioning and fission, but the molecular mechanisms controlling these processes remain poorly understood. Here, we identify a previously uncharacterized *P. falciparum* protein (Pf3D7_0613600), which we name PfAnchor, as a key regulator of apicoplast fission. Using Ultrastructure Expansion Microscopy (U-ExM), we show that PfAnchor localizes to the apicoplast throughout the asexual blood-stage. Conditional depletion disrupts apicoplast fission, leading to incomplete cytokinesis and parasite death. Notably, loss of the apicoplast’s elongated branched structure via azithromycin treatment rescues these defects, underscoring Anchor’s specific role in apicoplast fission. Immunoprecipitation identified an interaction with the dynamin-like GTPase PfDyn2, a key mediator of both apicoplast and mitochondrial fission, establishing PfAnchor as the first apicoplast-specific dynamin adaptor protein. Our findings define PfAnchor as an essential factor for apicoplast fission and inheritance in *P. falciparum* blood-stage parasites, highlighting parasite-specific organelle division as a potential vulnerability for therapeutic intervention.

## Introduction

Protists, a diverse group of predominantly single-celled eukaryotes, represent the largest component of eukaryotic diversity and are found in nearly every environment [1, 2]. Like other eukaryotes, most protists contain mitochondria and/or plastid-derived organelles that support essential metabolic processes. Examples include photosynthetic plastids in dinoflagellates and the non-photosynthetic apicoplast found in most apicomplexan parasites [3]. Both mitochondria and plastids originate from endosymbiotic events where ancestral eukaryotes engulfed bacteria or algae that evolved into organelles [4]. Evidence for this evolutionary origin includes the presence of organelle genomes within mitochondria and plastids, as well as the multi-membrane structure of plastids derived from secondary endosymbiosis [5]. Given their fundamental roles in survival and metabolism and the absence of plastids in human hosts, these organelles represent attractive targets for novel antiparasitic drug development [6, 7].

Malaria is a parasitic disease caused by protozoan *Plasmodium* parasites, with an estimated 249 million cases and 608,000 deaths reported in 2022 [8]. *Plasmodium* parasites invade host red blood cells and undergo multiple rounds of asynchronous DNA replication within a shared cytoplasm, resulting in a multi-nucleated parasite [9]. These parasites contain a non-photosynthetic plastid known as the apicoplast. In apicomplexans, the apicoplast derives from a red algal secondary endosymbiont, acquired before the divergence of dinoflagellates and apicomplexans. This is reflected in their four surrounding membranes and the SELMA (symbiont-specific ER-like machinery) protein import system, which are typical features of complex plastids of red algal origin [10–16]. The apicoplast has its own small genome of about 35 kb. While it aids in producing fatty acids and heme, its primary functions include synthesizing isoprenoid precursors such as isopentenyl pyrophosphate (IPP) and coenzyme A (CoA) during asexual blood stage development [17–23].

The apicoplast in *Plasmodium* is a single, branching organelle that expands in volume as the parasite progresses through the intraerythrocytic cycle. Volume electron microscopy (EM) studies, along with our recent work using ultrastructure expansion microscopy (U-ExM), have confirmed that the apicoplast undergoes multiple asynchronous fission events during schizogony, coinciding with parasite cytokinesis [24–26]. Recent studies have shown that apicoplast fission is mediated by a dynamin-like protein, PfDyn2 (Pf3D7_1037500), and is essential for parasite survival, as the apicoplast cannot be synthesized de novo [27, 28]. Unlike classical dynamin proteins, which contain a pleckstrin homology (PH) domain for direct membrane interaction during fission, dynamin-related proteins rely on adaptor proteins to recruit them to membranes and assemble functional fission complexes [29–31]. Mitochondrial fission adaptors vary widely in sequence and structure, with four identified in humans [32–35], three in yeast [36–38], two in the related parasite *Toxoplasma gondii* [39, 40], and one in *Plasmodium* [41]. However, no adaptor proteins specific to apicoplast fission have been identified in either *T. gondii* or *P. falciparum*, leaving the mechanism by which PfDyn2 facilitates apicoplast division unresolved.

In this study, we identify and characterize PfAnchor (Pf3D7_0613600), a nuclear-encoded protein that localizes to the apicoplast during the blood stage developmental cycle. Using the inducible TetR-DOZI conditional knockdown system [42], we demonstrate that PfAnchor is essential for parasite growth. Live microscopy of PfAnchor-deficient parasites revealed that while egress occurs, daughter cells form large clumps, preventing successful invasion. U-ExM analysis further showed that the apicoplast fails to undergo proper fission, resulting in most daughter parasites remaining tethered to each other via a collapsed apicoplast, while a subset of parasites emerges without an apicoplast. Blocking apicoplast branching via azithromycin treatment fully rescued the growth defect in PfAnchor-deficient parasites. Immunoprecipitation experiments identified an interaction between PfAnchor and PfDyn2, suggesting that PfAnchor acts as a dynamin adaptor protein required for apicoplast fission and inheritance. By uncovering a key regulator of apicoplast division, this study provides new insights into the molecular machinery governing organelle inheritance. Additionally, we highlight the potential of targeting apicoplast morphology as a novel strategy for inducing same-cycle parasite death, positioning PfAnchor as a promising therapeutic target for malaria intervention.

## Results

### PfAnchor: A conserved *Plasmodium* apicoplast-associated protein in the blood stage

Proximal labeling experiments using *P. falciparum* Minichromosome Maintenance-binding protein (PfMCMBP) as bait [43] led to the identification of an uncharacterized protein that we named PfAnchor (Pf3D7_0613600, **Figure S1-S2, Supp Data 1**). PfAnchor is a *Plasmodium specific* 1034 amino acid protein. No homologs of PfAnchor could be identified in other apicomplexan organisms, suggesting it may represent a lineage-specific adaptation. The sequence structure contains a hydrophobic helix at the N-terminal end, predicted to be a transmembrane domain, as well as a reported phosphorylation site at S204 identified in the schizont stage phosphoproteome of *P. falciparum* [44] (**Figure 1a**). The functional significance of this phosphorylation remains unknown and will require future investigation. Alphafold3 structure prediction [45] revealed an ordered region at the C-terminal end which is present across *Plasmodium* species (**Figure 1a, Figure S3**). By combining Alphafold3 predictions with a DALI search [46], we, along with a recent publication [47], identified this region as a putative pleckstrin homology (PH) domain (**Figure 1a**). These domains are typically involved in membrane binding suggesting that PfAnchor may associate with a membrane. Sequence analysis of the predicted PH domain revealed strong conservation among *Plasmodium* species, with the highest similarity between *P. falciparum* and the *Laverania* subgenus (**Figure S3a**). Alphafold3 structure predictions across *Plasmodium* species with varying degrees of PfAnchor homology showed structural conservation of the predicted PH domain with high confidence model prediction in multiple species (**Figure S3b**). This structural conversation suggests that the predicted PH domain may have a conserved functional role in *Plasmodium* parasites.

**Figure 1:**
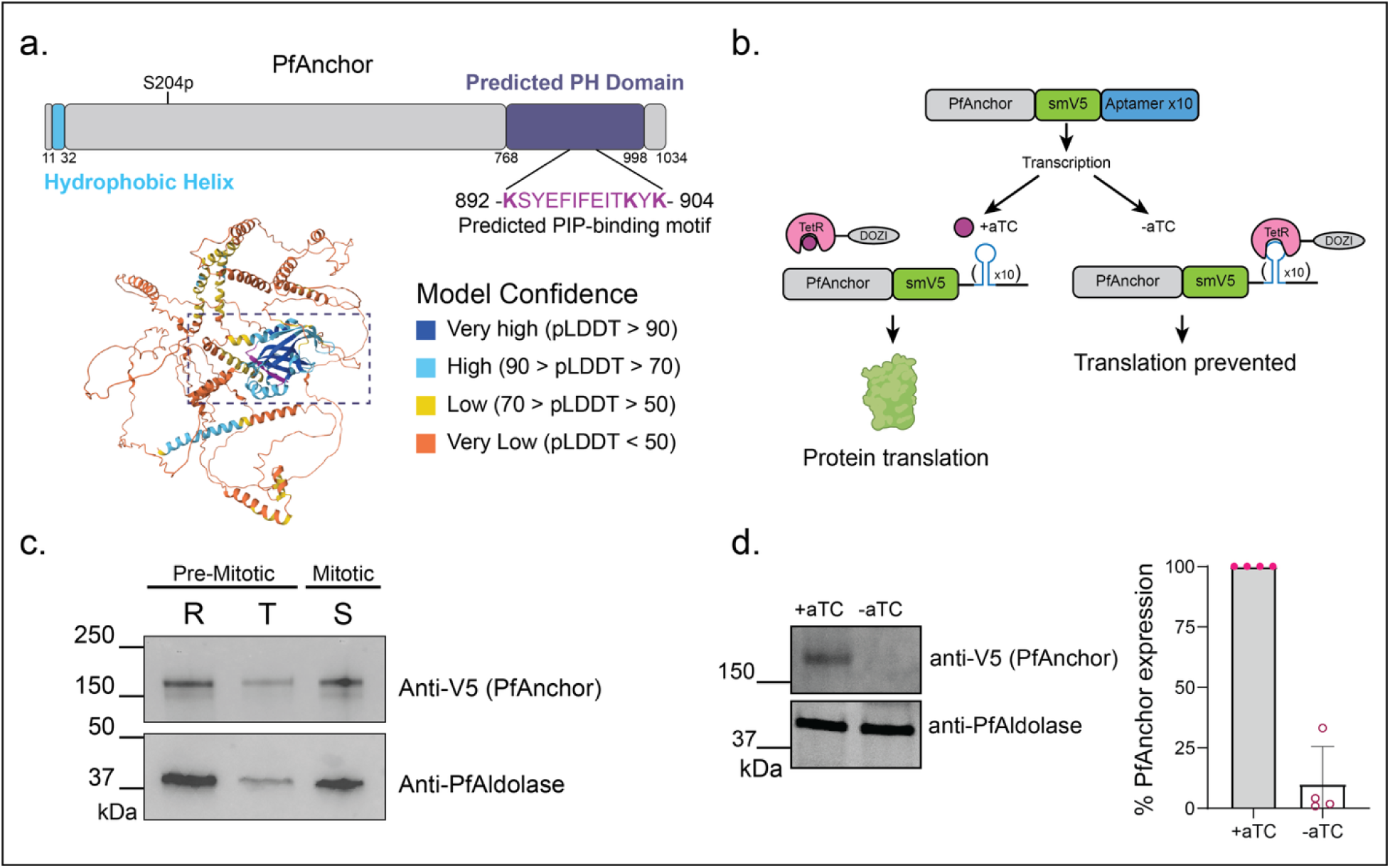
PfAnchor is an essential conserved *Plasmodium* protein expressed through the asexual life cycle. **a.** Schematic representation of PfAnchor’s structural features and predicted protein folding. PfAnchor primary sequence shows a single phosphorylation site at S204. Alphafold3 modeling combined with a DALI protein domain search identified a conserved pleckstrin homology (PH) domain (PfAnchor AA 768-998) across *Plasmodium* species. The sequence alignment of this region shows strong conservation, with a predicted phosphoinositides (PIP)-binding motif (PfAnchor AA 892-904) highlighted in magenta in the structural model. **b.** Schematic of PfAnchor tagging strategy and conditional knockdown system. PfAnchor was tagged with an smV5 epitope for visualization, and knockdown was achieved using the TetR-aptamer system, where protein translation depends on anhydrotetracycline (aTC) availability. **c.** Representative western blot (N=3) of parasite protein lysates from ring (R), trophozoite (T), and schizont (S) stage. Anti-V5 detects PfAnchor, showing that it is expressed throughout the asexual life cycle with highest abundance in schizont stages. Anti-PfAldolase is used as a loading control, as it is a constitutively expressed cytoplasmic protein that maintains relatively stable levels across all intraerythrocytic stages. **d.** Western blot analysis confirming PfAnchor knockdown efficiency. Quantification performed via measurement of the fluorescent intensities of the sample (V5) bands compared to the normalized intensities of the loading control (PfAldolase) bands. Plotted is mean +/- SD, n = four biological replicates.

Having identified PfAnchor as a potential membrane-associated protein through structural predictions, we next investigated its subcellular localization and membrane association. To enable detection and conditional regulation of PfAnchor, we engineered a C-terminal fusion of the protein with the spaghetti monster-V5 (smV5) epitope tag [48] and integrated the TetR-based conditional knockdown system [42] into the 3D7 parasite strain, named PfAnchor iKD (**Figure 1b, Figure S4**). This system allows for controlled expression of PfAnchor in the presence of anhydrotetracycline (aTC), a small molecule that stabilizes mRNA translation. In the absence of aTC, translation is suppressed, resulting in effective protein knockdown. Consistent with PfAnchor’s transcriptional profile [49] we were able to detect protein expression throughout the asexual blood stages of the parasite life cycle (**Figure 1c**). To validate the functionality of the inducible knockdown system, we washed out aTC one cycle prior to analysis, approximately 10 hours before egress, and collected samples 40 hours post-invasion. Western blot analysis showed that PfAnchor expression was reduced by more than 90% across four independent biological replicates, confirming the efficacy of the knockdown system (**Figure 1d**).

To test whether PfAnchor associates with membranes, we performed sodium carbonate solubility assays. Parasites were hypotonically lysed to remove soluble cytosolic proteins, then sequentially extracted with sodium carbonate (to release peripherally associated membrane proteins) and Triton X-100 (to solubilize integral membrane proteins). The supernatant and pellet fractions were analyzed by western blot. As controls, we probed our blots for GFP (non-membrane associated PfACP-GFP), PfAldolase (cytosolic and peripherally associated) and PfEXP2 (integral membrane). PfAnchor was not detected in the hypotonic fraction, indicating it is not soluble in the cytosol and instead associates with membranes. However, because we lack a definitive peripheral membrane control, we cannot confidently determine whether PfAnchor is a peripheral or integral membrane protein **(Figure S5**).

Given its membrane association, we next sought to identify the specific membrane compartment with which PfAnchor associates. Using U-ExM, we observed that PfAnchor localizes to regions containing non-nuclear DNA, suggesting a potential association with a DNA-containing organelle. By staining PfAnchor parasites with MitoTracker, we showed that PfAnchor did not co-localize with the mitochondrion (**Figure 2a, b**). As the apicoplast possesses its own genome, we hypothesized that PfAnchor might localize to the apicoplast during this stage. To test this, we generated a new cell line by tagging PfAnchor with smV5 in the PfMev::Api-GFP background [50] that we named PfAnchor^Mev^ iKD (**Figure S5**). This strain enables conditional disruption of the apicoplast by mevalonate (Mev) supplementation, supporting viability of the parasite without the organelle. This system facilitates visualization of the apicoplast through GFP tagging and allows investigation of PfAnchor’s function in the absence of an intact apicoplast. Imaging analysis of this cell line revealed that PfAnchor localizes to the periphery of the apicoplast throughout schizogony (**Figure 2b**), supporting its involvement in apicoplast biology.

**Figure 2:**
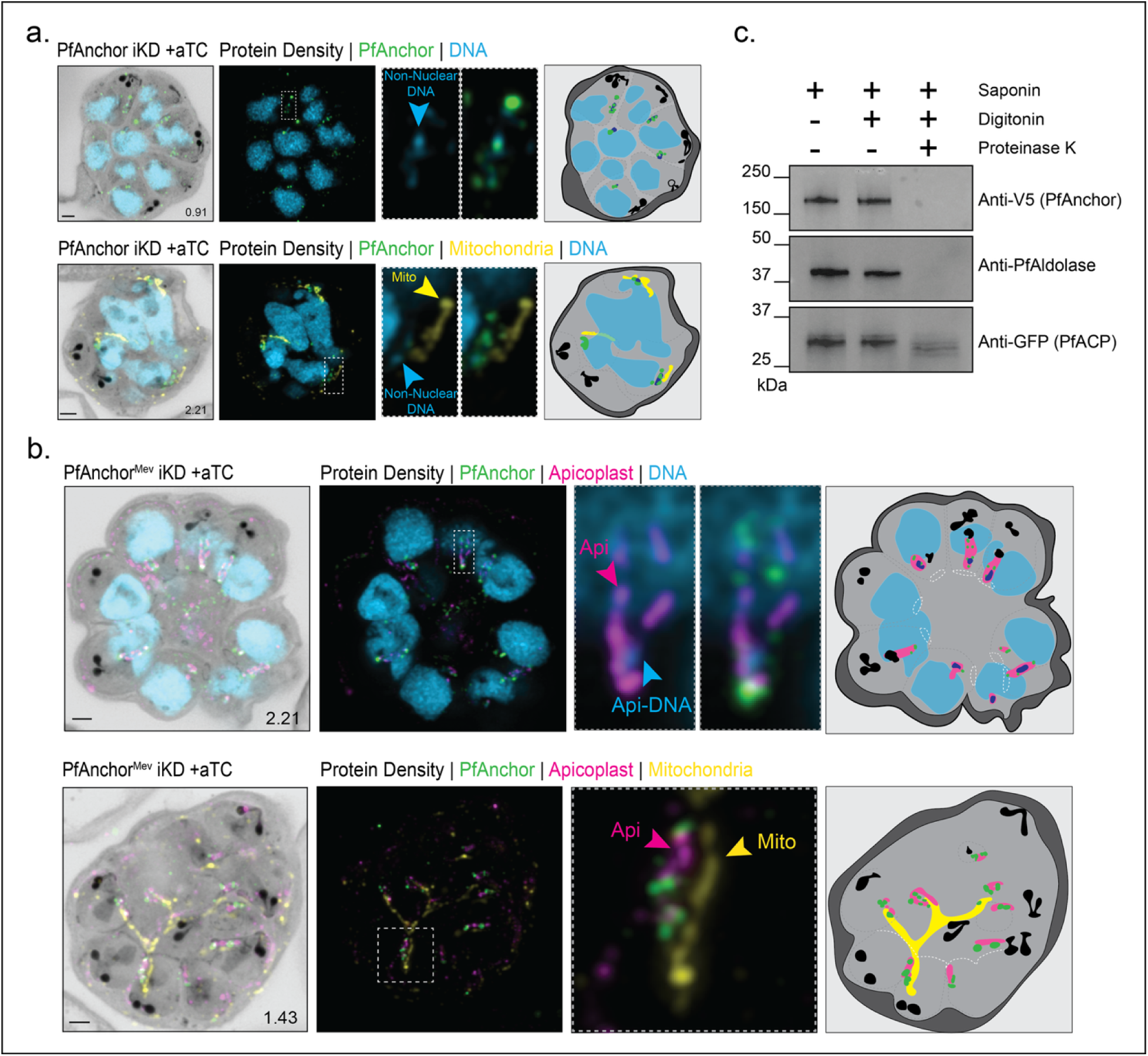
PfAnchor localizes to the cytoplasmic side of the apicoplast outer membrane. **a-b.** U-EXM analysis of PfAnchor localization in PfAnchor-iKD and PfAnchor^Mev^-iKD cell lines. PfAnchor is detected in close proximity to non-nuclear DNA but does not colocalize with mitochondrial DNA. Apicoplast staining confirms that PfAnchor is specifically localized around the apicoplast. Protein density is shown in grayscale, PfAnchor in green, apicoplast in magenta, mitochondria in yellow, and DNA in blue. Scale bars: 2 µm, with image depth (µm) indicated. Schematic representations highlight non-nuclear DNA (dark blue) and basal complexes (white), with light gray dashed lines indicating areas of potential membrane staining based on changes in NHS ester density. **c.** The proteinase K protection assay indicates that the C-terminal tagged PfAnchor is cytoplasmically exposed. Anti-PfAldolase and anti-GFP (PfACP) were used as unprotected and protected controls, respectively.

To determine the orientation of PfAnchor within the apicoplast, we performed proteinase K protection assays. Infected red blood cells were treated with saponin to selectively permeabilize the host cell membrane and release hemoglobin, while leaving the parasites enclosed within the parasitophorous vacuole membrane and the host RBC membrane. Then treated with digitonin to selectively permeabilize the parasite plasma membrane. The parasite pellets were subsequently exposed to proteinase K, which degrades cytosolic-facing proteins while leaving proteins within organellar compartments intact. We used PfAldolase as a control for cytosolic and cytosolic-facing proteins, and GFP as a control for protected proteins, since our engineered parasites expressed PfACP-GFP, a luminal apicoplast protein. PfAnchor was sensitive to proteinase K digestion, indicating that its C-terminal epitope tag is exposed to the cytosol (**Figure 2c**). In contrast, GFP remained detectable, confirming that proteins inside the apicoplast lumen are shielded from protease treatment. These results together confirm that PfAnchor is a cytosol-facing protein associated with the apicoplast membrane.

### PfAnchor is essential for apicoplast partitioning and merozoite formation

To investigate the role of PfAnchor in parasite development, we removed anhydrotetracycline (aTC) from the culture and monitored parasite growth by microscopy. PfAnchor-deficient parasites exhibited a complete growth arrest during the first cycle of depletion, with no early-stage parasites detected in the culture, suggesting a critical role in either egress or invasion (**Figure 3a**). To test this, we performed live microscopy on PfAnchor-expressing and PfAnchor-deficient parasites to investigate whether egress was affected. Parasites were treated with Compound 1 to arrest and synchronize them just prior to egress from red blood cells [51]. After washing with fresh media, egress events were monitored. PfAnchor-expressing parasites displayed rapid and explosive egress, with individual daughter merozoites dispersing quickly. By contrast, PfAnchor-deficient parasites underwent egress, yet the majority of daughter merozoites remained clustered and failed to fully separate, often remaining attached near the food vacuole (**Figure 3b, c**). To quantify egress phenotypes, we categorized parasites based on morphology: “normal” morphology was defined as minimal merozoite clumping with daughter parasites dispersing effectively, while “abnormal” morphology was characterized by limited separation of daughter parasites, which formed slow-moving or immobile clumps. PfAnchor-expressing parasites showed an average of 89.3 +/- 9.5% normal morphology (52/61 egress events) over three biological replicates while PfAnchor-deficient parasites showed an average of 10.7 +/- 9.3% normal morphology (3/39 egress events) over three biological replicates. We note that PfAnchor-deficient parasite morphology closely resembles the phenotype associated with impaired apicoplast segregation [52]. Given this resemblance, we hypothesized that disrupted apicoplast division may underlie the observed defect in cytokinesis. To test this, we visualized the apicoplast morphology in PfAnchor-deficient parasites expressing ACP-GFP (PfAnchor^Mev^ iKD parasites). Imaging PfAnchor-deficient parasites revealed two distinct groups of merozoites: clumped merozoites that share a single, unsegregated apicoplast, and free merozoites that display no apicoplast signal. In contrast, PfAnchor-expressing parasites exhibit proper segregation, with each merozoite containing a distinct apicoplast (**Figure 3d**). These findings demonstrate that PfAnchor is essential for apicoplast inheritance during merozoite formation.

**Figure 3:**
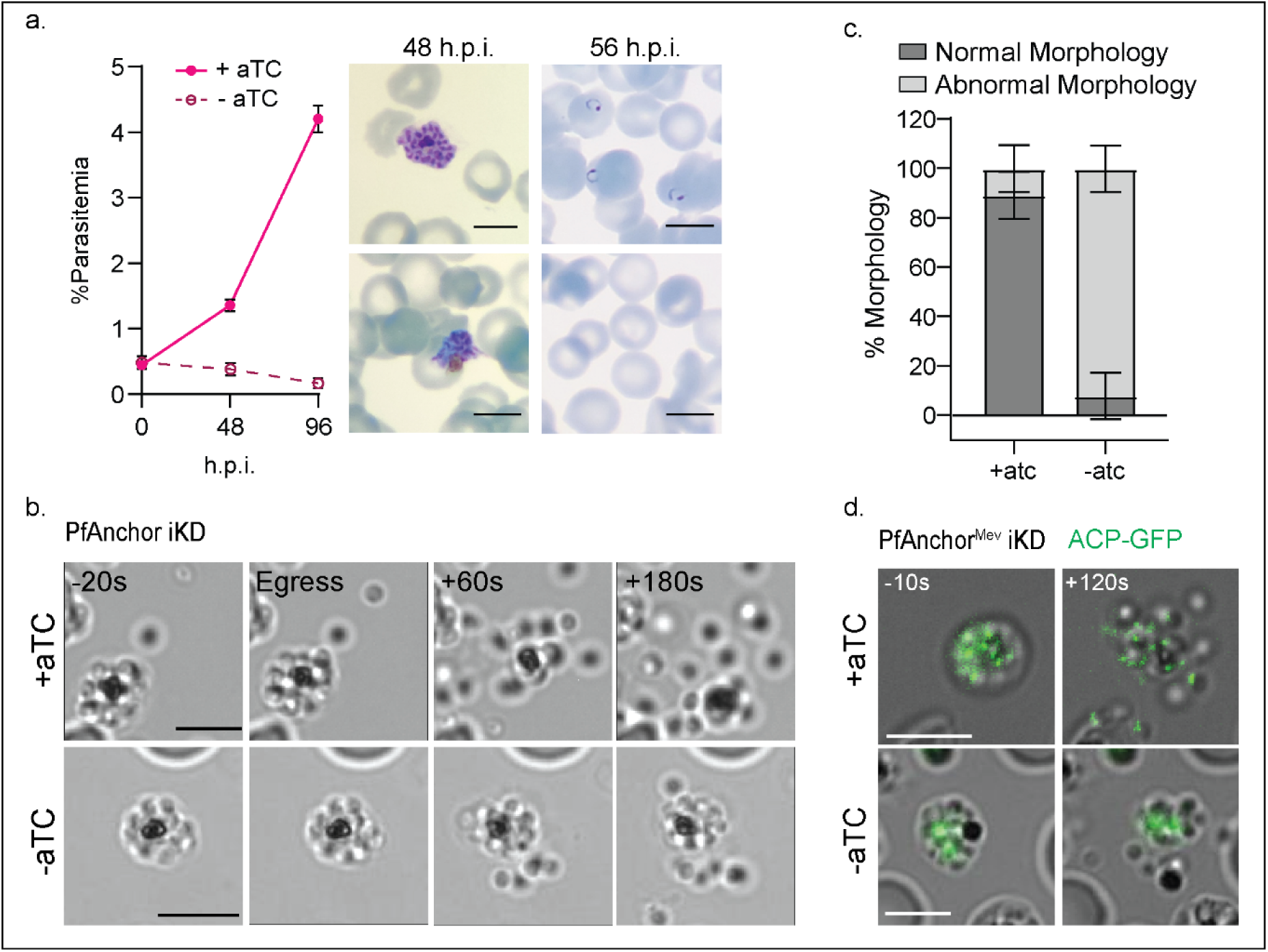
PfAnchor is required for the release of segregated merozoites but dispensable for egress. **a.** Growth analysis of PfAnchor-expressing and deficient parasites over two replication cycles. Parasitemia was quantified by light microscopy with three biological replicates. Hemacolor-stained smears show abnormal late-stage morphology prior to egress and no ring-stage parasites after egress in PfAnchor-deficient cultures. Scale bar 5 µm. **b.** Representative live microscopy images of PfAnchor-expressing (top) and PfAnchor-deficient (bottom) parasites at egress. While PfAnchor-expressing parasites release individual merozoites, PfAnchor-deficient parasites form clumps of daughter cells that remain tethered (abnormal morphology). Scale bar 5 µm. **c.** Quantification of normal and abnormal daughter parasite morphologies after egress from live microscopy experiments over three biological replicates. PfAnchor-expressing parasites had 89.3 ± 9.5% normal morphology across three replicates, whereas PfAnchor-deficient parasites showed 10.7 ± 9.3% normal morphology. **d.** Representative live microscopy images of PfAnchor-expressing (top) and PfAnchor-deficient (bottom) parasites at egress, with the apicoplast labeled in green. Parasites expressing PfAnchor show daughters moving apart with the apicoplast labeled in green (ACP-GFP). In PfAnchor-expressing parasites, daughter cells separate and properly inherit apicoplast fragments. In contrast, in PfAnchor-deficient parasites, daughters remain clumped with persistent apicoplast material across the cluster or retained in the residual body. Scale bar 5 µm.

### PfAnchor is required for apicoplast division during IDC

Given its localization to the apicoplast periphery and the observed defects in organelle inheritance upon knockdown, we reasoned that PfAnchor plays a direct role in apicoplast division. We hypothesize that in the absence of PfAnchor, apicoplast fission fails, resulting in merozoites that are segregated but remain tethered by a single, undivided apicoplast. To test this, we used U-ExM to track apicoplast division during parasite replication and determine how PfAnchor depletion disrupts organelle division and inheritance. To assess whether PfAnchor loss affects apicoplast biogenesis, we measured apicoplast area in cell projections [25]. No significant differences were observed between PfAnchor-expressing and deficient parasites, indicating that apicoplast biogenesis remains unaffected (**Figure 4a**). Additionally, as we and others previously demonstrated, the apicoplast adopts a characteristic crown shape during early segmentation, with a single branched apicoplast connecting all outer centriolar plaques (CPs) [25, 53, 54]. This organization was also observed in PfAnchor-deficient parasites, indicating that the pre-fission positioning of the apicoplast and its association with the CPs remain unaffected in the absence of PfAnchor (**Figure S6**). However, PfAnchor-deficient parasites exhibited a striking defect in apicoplast inheritance during cytokinesis. Unlike control parasites, which display individual, segmented apicoplasts in their daughter parasites, PfAnchor-deficient parasites retained a single, interconnected apicoplast that linked multiple daughter parasites (**Figure 4b, c**). Live microscopy revealed that egress occurred at normal timing; however, instead of releasing individual merozoites, PfAnchor-deficient parasites produced clusters of daughters that remained interconnected by undivided apicoplast material (**Figure 3**). U-ExM further confirmed this phenotype, showing that these apicoplast-negative merozoites had separated from the cluster and moved freely (**Figure 4d**). This resulted in two distinct populations of daughter parasites in the PfAnchor-deficient samples: (1) parasites connected by a shared apicoplast traversing the basal complex (**Figure 4e**), and (2) parasites that were physically separated from the residual body but devoid of apicoplast signal (**Figure 4d**). Notably, while PfAnchor-deficient parasites displayed severe apicoplast inheritance defects, mitochondria biogenesis and division remained unaffected, with each daughter cell inheriting a single mitochondrion (**Figure S7**). These findings establish PfAnchor as a critical factor of apicoplast division and inheritance, and its loss likely prevents parasite propagation and survival.

**Figure 4:**
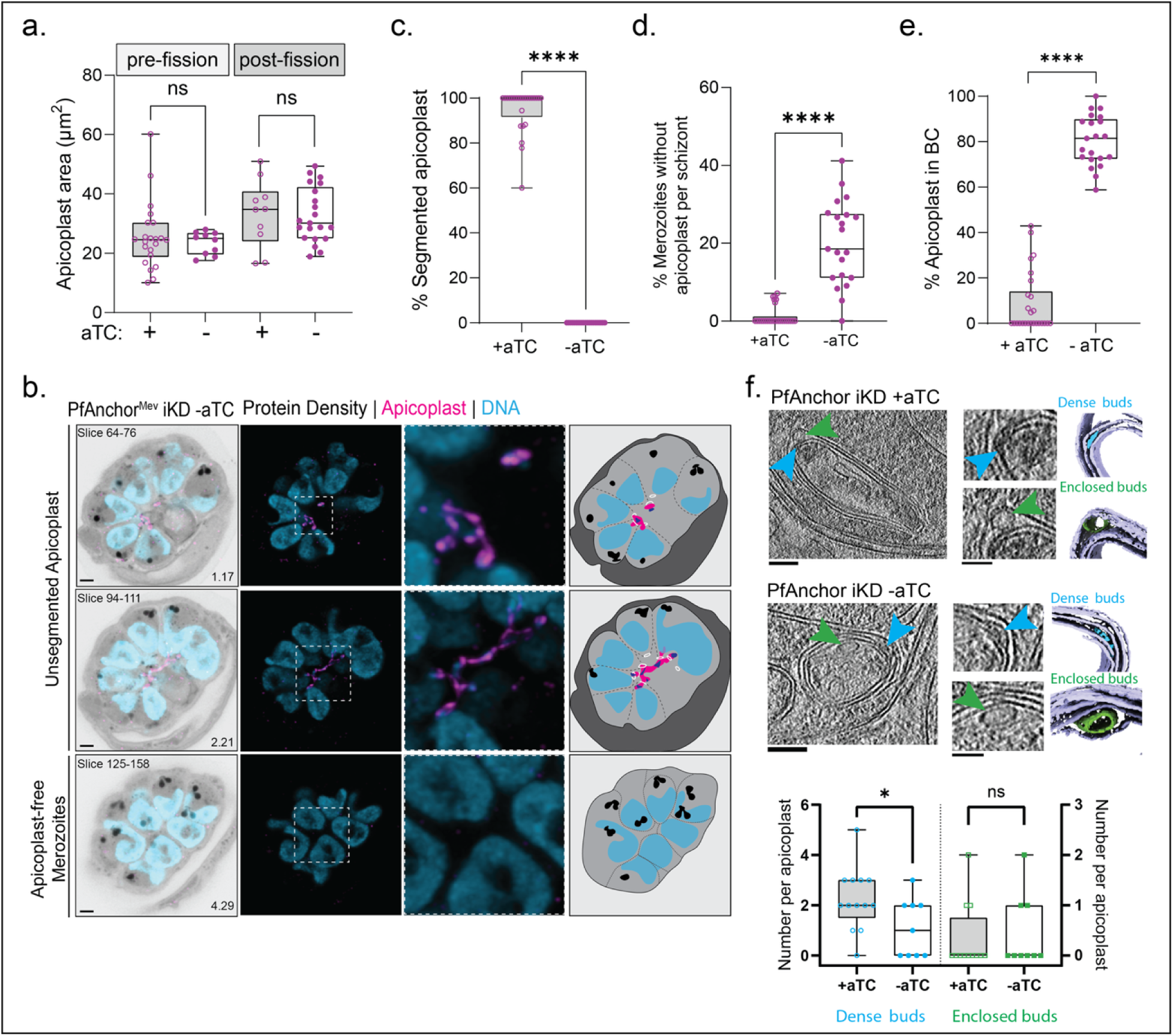
PfAnchor is required for apicoplast division and inheritance but not biogenesis. **a.** Quantification of total apicoplast area in age-matched pre- and post-division parasites shows no significant differences between PfAnchor-expressing and PfAnchor-deficient parasites, indicating that apicoplast biogenesis is unaffected (unpaired t-tests, +aTC: 32 cells over 2 replicates, -aTC 31 cells over 2 replicates, not significant). **b.** U_ExM representative images illustrating the quantified phenotype in (c-e). Shown are z-slices illustrating (top) apicoplast signal trapped in the basal complex preventing final separation of daughter cells, (middle) an unsegmented apicoplast spanning daughters, and (bottom) apicoplast-free merozoites. Protein density in grayscale, apicoplast in pink and DNA in blue. Scale bar 2 µm. Z-slices used are indicated, with image depth (µm) noted. **c.** Percentage of daughter parasites per schizont with a fully segmented apicoplast in schizonts stalled at the end of segmentation. Unpaired t-tests, +aTC: 25 cells over 3 replicates, -aTC: 21 cells over 3 replicates, **** indicates p-value < 0.0001. **d.** Percentage of daughter parasites lacking apicoplast signal in schizonts stalled at the end of schizogony. Unpaired t-tests, +aTC: 25 cells over 3 replicates, -aTC: 21 cells over 3 replicates, **** indicates p-value < 0.0001. **e.** Percentage of daughter parasites per schizont exhibiting apicoplast signal within the basal complex (BC) in schizonts stalled at the end of schizogony. Unpaired t-tests, +aTC: 25 cells over 3 replicates, -aTC: 21 cells over 3 replicates, **** indicates p-value < 0.0001. **f.** Representative cryo-electron tomography (cryo-ET) images showing that PfAnchor depletion does not affect apicoplast membrane integrity. However, analysis of internal apicoplast structures reveals a significant reduction in the number of dense intermembrane buds (blue arrows and data points) upon PfAnchor knockdown, while enclosed membrane buds (green arrows and data points) remain unaffected. Scale bars: 100 nm (unpaired t-tests, * indicates p-value < 0.05).

To further investigate the impact of Anchor depletion on apicoplast structure, we performed cryo-focused ion beam (cryo-FIB) milling and cryo-electron tomography (cryo-ET) on PfAnchor iKD parasites in the presence and absence of aTC. To ensure stage-matched analysis, we treated cultures with Compound 1 (4-[2-(4-fluorophenyl)-5(1-methylpiperidine-4-yl)-1H-pyrrol-3-yl]pyridine or C1), a reversible protein kinase G inhibitor that arrests schizonts immediately prior to egress by preventing parasitophorous vacuole membrane (PVM) rupture [51, 55]. C1-stalled schizonts were Percoll-purified and incubated in media containing 2.5 µM C1 for 2–3 hours before fixation. We analyzed a total of 12 tomograms from PfAnchor-expressing (+aTC) parasites, all containing identifiable apicoplasts, and 9 tomograms from PfAnchor-deficient (–aTC) parasites that featured apicoplasts. The apicoplast was distinguished in tomograms based on its characteristic four-membrane structure. 2D Measurement of the smallest internal diameter, taken from a 2D slice at the center of the apicoplast volume, revealed no significant difference between PfAnchor-expressing and PfAnchor-deficient parasites (**Figure S8a**). In both PfAnchor-expressing and PfAnchor-deficient parasites, we observed cryo-EM densities within the intermembrane space of the apicoplast (**Figure 4f**). These densities included dense intermembrane buds, defined as electron-dense tethering structures between neighboring apicoplast membranes (**Figure 4f, blue arrows**). Interestingly, the frequency of these dense intermembrane buds was significantly reduced in PfAnchor-deficient parasites, with PfAnchor-expressing parasites present ∼2.2 dense buds per apicoplast on average (S.D. = 1.2), while PfAnchor-deficient parasites averaged ∼1.1 dense buds per apicoplast (S.D. = 1.2).

In addition to the dense intermembrane buds, we identified enclosed membrane-like buds, small, circular densities forming between apicoplast membranes (**Figure 4f, green arrows**). Unlike the dense intermembrane buds, the number of enclosed buds remained unchanged between PfAnchor-expressing and PfAnchor-deficient parasites. This suggests that while PfAnchor influences the formation or maintenance of dense intermembrane buds, it does not significantly impact the presence of enclosed buds, indicating that some aspects of membrane organization remain intact. Beyond these structures, we observed additional internal features within the apicoplast lumen. These included single-layer and double-layer membrane structures, which may represent distinct internal compartments or intermediates in apicoplast remodeling (**Figure S8b-e**). Quantification of single-layer membrane structures revealed no significant differences in their abundance or internal diameter between PfAnchor-expressing and PfAnchor-deficient parasites (**Figure S8c, d**), suggesting that their formation is independent of PfAnchor. However, double-layer membrane structures were exclusively detected in PfAnchor-deficient parasites (**Figure S8e**), suggesting that PfAnchor depletion alters the internal architecture of the apicoplast. One important note is that the parasites imaged in this study were chemically fixed, a method we used to preserve cellular structures in the absence of local cryo-preservation capabilities. However, we also examined cryo-preserved *Plasmodium* tomograms [56] and similarly observed the presence of dense buds. This comparison with unfixed samples (**Figure S9**) confirms that the dense buds are not artifacts of chemical fixation. These findings suggest that while PfAnchor is not required for overall membrane biogenesis, it may play a role in maintaining the organization or remodeling of the apicoplast membrane, potentially influencing its division and inheritance.

### Loss of apicoplast-branched structure rescues growth in PfAnchor-deficient parasites

We hypothesized that the egress defects observed in PfAnchor-deficient parasites result from physical tethering of daughter merozoites by the branched apicoplast structure and that disrupting this tethering could restore parasite viability. Bacterial translation inhibitors such as doxycycline, clindamycin, and azithromycin are known to disrupt apicoplast structure, leading to the formation of apicoplast-like vesicles during the second cycle of treatment and parasite death [17, 50]. The growth of parasites treated with bacterial translation inhibitors can be rescued with the addition of exogenous IPP [17], leading to parasites growing with apicoplast-like vesicles instead of a branching apicoplast and irreversible loss of the apicoplast genome [57]. The high cost and the difficulties associated with the IPP-rescue system [17] for long-term studies led to the development of an apicoplast bypass system by Swift et al. in 2020 [50]. This system introduces four enzymes into the cytoplasm of the parasite, allowing for IPP synthesis via the mevalonate (Mev) pathway in the cytoplasm, thereby bypassing the endogenous methylerythritol phosphate (MEP) pathway in the apicoplast. By adding mevalonate to the growth medium, the bypass system significantly reduces the cost and improves the feasibility of long-term experimental studies.

To test our hypothesis, we used the apicoplast bypass PfAnchor^Mev^ iKD cell line and treated parasites with the antibiotic azithromycin (AZ) to prevent apicoplast branching during PfAnchor knockdown [17, 58]. Briefly, PfAnchor^Mev^ iKD parasites were initially cultured in the presence of 50 µM mevalonate (Mev) and 100 nM AZ for 7 days. The parasites were then split into four experimental groups: (1) +aTC -Mev (green), (2) -aTC -Mev (orange), (3) +aTC +Mev (blue), and (4) -aTC +Mev (red) (**Figure 5a**). These groups were cultured for an additional 8 days, taking samples every 2 days for growth analysis. To prevent culture collapse, parasite cultures were diluted (1:8) every 2 days. U-ExM validated the loss of the apicoplast branching structure and PCR analysis confirmed the loss of the apicoplast genome regardless of the presence of PfAnchor (**Figure 5c, g, Figure S10**).

**Figure 5:**
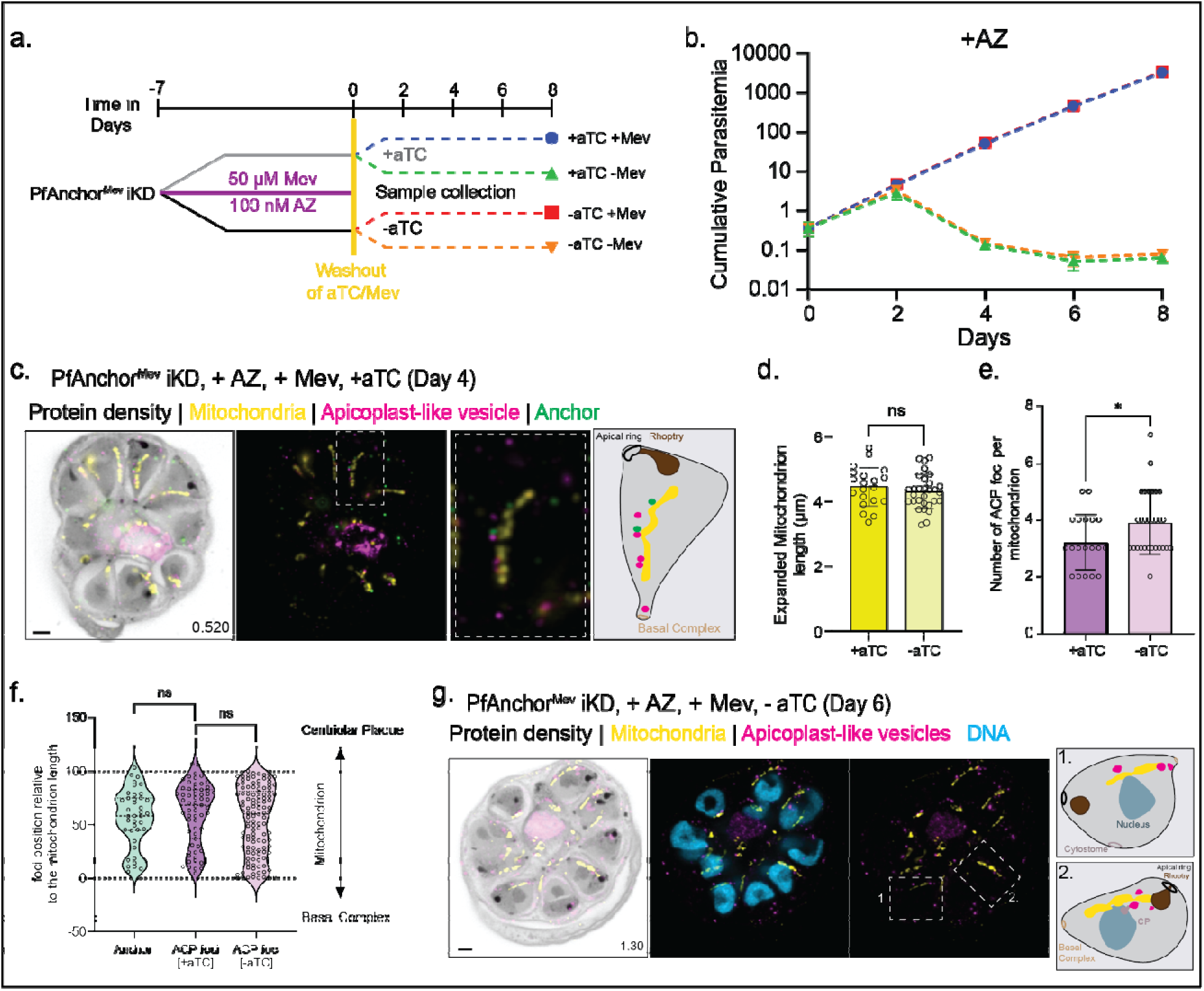
Disruption of apicoplast branching rescues PfAnchor-deficient parasites, while apicoplast-like vesicle distribution along mitochondria remain unchanged. **a.** Schematic of the experimental setup. PfAnchor^Mev^ iKD parasites were pre-treated with mevalonate (+Mev) and azithromycin (+AZ) for 7 days to disrupt the apicoplast structure, leading to the formation of apicoplast-like vesicles. Following this pre-treatment, parasites were washed and maintained under the indicated conditions (±aTC, ±Mev) for downstream analyses. **b.** Cumulative parasitemia of PfAnchor-expressing (+aTC) and PfAnchor-deficient (-aTC) parasites over 8 days in +AZ conditions. In AZ-treated parasites, disruption of the apicoplast resulted in the formation of apicoplast-like vesicles, which rescued the growth defect of PfAnchor-deficient parasites, but only in the presence of mevalonate (+Mev). Parasites lacking both PfAnchor and mevalonate (-Mev) failed to propagate. Parasites were sampled every 2 days for growth and PCR analysis, with cultures diluted at 1:8 every 2 days to ensure continued replication. Data represent mean ± SD from 2 biological replicates in quadruplicate. **C**. Representative U-ExM image of a PfAnchor-expressing parasite displaying apicoplast-like vesicles (pink, anti-GFP) within the forming daughter parasites, along with PfAnchor (green, anti-V5) and mitochondria (yellow, anti-Hsp60). Protein density is shown in grayscale. Scale bar: 2 µm, with z-slices and image depth (µm) indicated. **d.** Quantification of mitochondrial length in PfAnchor-expressing (8 cells, 15 mitochondria) and PfAnchor-deficient (10 cells, 30 mitochondria) parasites following AZ treatment. No significant differences in mitochondrial length were observed between conditions (ns, unpaired t-test). (**e-g**). Apicoplast-like vesicles were analyzed for their association with mitochondria. **e**. The number of apicoplast-like vesicles associated with mitochondria was significantly increased in PfAnchor-deficient (-aTC) parasites compared to PfAnchor-expressing (+aTC) parasites. **f**. Violin plots show the spatial distribution of apicoplast-like vesicles along the mitochondria, demonstrating that PfAnchor depletion does not alter their positioning. Statistical analysis was performed using unpaired t-tests, +aTC: 20 mitochondria from 8 schizonts; -aTC: 30 mitochondria from 15 schizonts. **g**. Representative U-ExM image of a PfAnchor-deficient parasite showing apicoplast-like vesicles (pink, anti-GFP) within forming daughter parasites, along with mitochondria (yellow, anti-Hsp60). Protein density is shown in grayscale. Scale bar: 2 µm, with z-slices and image depth (µm) indicated.

Growth analysis revealed distinct phenotypes depending on PfAnchor expression and the presence of Mev. In parasites with apicoplast-like vesicles (+AZ), parasites supplemented with Mev displayed equivalent growth rates regardless of PfAnchor expression (**blue and red lines overlapping, Figure 5b**), indicating that disruption of the branched apicoplast structure fully rescued the growth defects associated with PfAnchor depletion. Conversely, parasites with apicoplast-like vesicles but not supplemented with Mev failed to grow regardless of PfAnchor expression (**green and orange lines overlapping, Figure 5b**), validating the experimental setup. These results demonstrate that the loss of the branched apicoplast structure is sufficient to fully rescue the growth defects in PfAnchor-deficient parasites. This finding underscores the critical role of PfAnchor in apicoplast fission. Since parasite viability is restored in the absence of a branched apicoplast, it appears unlikely that PfAnchor has any other essential functions beyond facilitating apicoplast fission.

As controls we tested the growth of parasites with a branched apicoplast (-AZ) in the presence or absence of Mev or PfAnchor. Branched apicoplast PfAnchor-expressing parasites showed robust growth with or without Mev supplementation (**green and blue lines overlapping, Figure S11**). Branched apicoplast PfAnchor-deficient parasites failed to grow without Mev, as evidenced by a decline in parasitemia (**orange line, Figure S11b**). Interestingly, PfAnchor-deficient parasites supplemented with Mev initially exhibited reduced growth but subsequently recovered, achieving growth rates comparable to PfAnchor-expressing parasites (**red line, Figure S11b**). We hypothesize that this initial recovery of growth is caused by free merozoites, which contain no apicoplast in the PfAnchor population (**Figures 3d**, **4b**) that can survive and propagate when supplemented with Mev. To test this hypothesis, we performed additional analysis on PfAnchor-deficient Mev-supplemented parasites at day 8 post aTC removal (**Figure S11c-e**). PCR results indicate that PfAnchor-deficient parasites don’t contain an apicoplast genome, while the nuclear and mitochondrial genomes remain intact. In addition, U-ExM imaging of parasites supplemented with Mev at day 8 post PfAnchor removal shows the same apicoplast-like vesicles that were formed after treatment with AZ (**Figure 5c, Figure S11e**), indicating loss of apicoplast structure and formation of vesicle-like structures.

### Apicoplast-like vesicles are retained in daughter parasites in association with mitochondria, independent of PfAnchor

To investigate the fate of the apicoplast following its disruption by azithromycin, we examined the spatial organization and inheritance of the resulting apicoplast-like vesicles using U-ExM (**Figure 5c, g**). In PfAnchor-expressing parasites, the majority of apicoplast-like vesicles were excluded from the daughter parasites and instead localized within the residual body at the completion of cytokinesis (**Figure S12**). However, a subset of apicoplast-like vesicles remained within daughter cells, and these retained vesicles were consistently associated with the mitochondrion (**Figure 5c, S12**). This suggests that mitochondrial association plays a key role in apicoplast vesicle inheritance, while vesicles not tethered to the mitochondrion are largely excluded from daughter parasites during cytokinesis.

To determine whether PfAnchor is required for apicoplast-like vesicle inheritance, we examined PfAnchor-deficient parasites and found that their vesicle localization and distribution were largely similar to PfAnchor-expressing parasites (**Figure 5g, S13**). However, quantification revealed a modest but significant increase in the number of mitochondrion-associated apicoplast vesicles in PfAnchor-deficient parasites compared to PfAnchor-expressing parasites (**Figure 5e**). This suggests that while PfAnchor is not required for vesicle retention, its depletion may subtly influence the balance between vesicle inheritance and clearance. Despite this increase, the spatial distribution of apicoplast-like vesicles along the mitochondrion was unchanged between conditions (**Figure 5f**), reinforcing the conclusion that PfAnchor does not impact the positioning of retained vesicles. As the mitochondrion served as a spatial reference for apicoplast vesicle distribution, we quantified mitochondrial length to ensure that mitochondrial organization remained unchanged in the presence or absence of PfAnchor. No significant differences in mitochondrial length were observed between PfAnchor-expressing and PfAnchor-deficient parasites following azithromycin treatment (**Figure 5d**), confirming that any differences in apicoplast-like vesicle localization would not be due to changes in mitochondrial morphology. Together, these data indicate that mitochondrial association is a key determinant of apicoplast vesicle retention, while PfAnchor is not required for their inheritance.

### PfAnchor interacts with PfDyn2 to facilitate apicoplast fission

Knockdown of PfAnchor disrupts apicoplast segregation and positioning, yet its precise molecular function remains unclear. To identify potential interacting partners, we performed immunoprecipitation (IP) experiments using two PfAnchor-tagged cell lines (PfAnchor iKD and PfAnchor^Mev^ iKD), followed by unbiased mass spectrometry. Parental untagged (3D7-Cas9) parasites served as a control to distinguish specific interactors from background proteins. Principal component analysis (PCA) of the mass spectrometry data confirmed the specificity of PfAnchor-associated interactions, as PfAnchor clustered distinctly from background interactors (**Figure 6a, Supp Data 2**). Among the most highly enriched proteins, PfDyn2 emerged as a primary interactor, supporting its role as a key regulator of apicoplast and mitochondrial fission [27, 28]. In addition to PfDyn2, PCA identified PfHsp70 (PF3D7_0818900) as a secondary interactor, suggesting a possible role in stabilizing protein complexes during apicoplast division [59, 60]. Beyond the PCA analysis, PfAnchor pulldown experiments identified additional interactors uniquely present in the PfAnchor dataset (**Figure 6c**). Among these, PfActin1 was specifically enriched, consistent with its established role in cytoskeletal regulation of organelle segregation [52, 61]. PfCinch, a basal complex protein [62], was also identified, suggesting a possible link between PfAnchor and cytoskeletal structures involved in organelle positioning. These findings provide strong evidence that PfAnchor interacts with PfDyn2, reinforcing its involvement in apicoplast fission. The identification of PfHsp70 as a secondary interactor suggests a potential chaperone function that may contribute to the stability of the fission complex. Additionally, the enrichment of PfActin1 and PfCinch supports a possible cytoskeletal role in apicoplast segregation.

**Figure 6:**
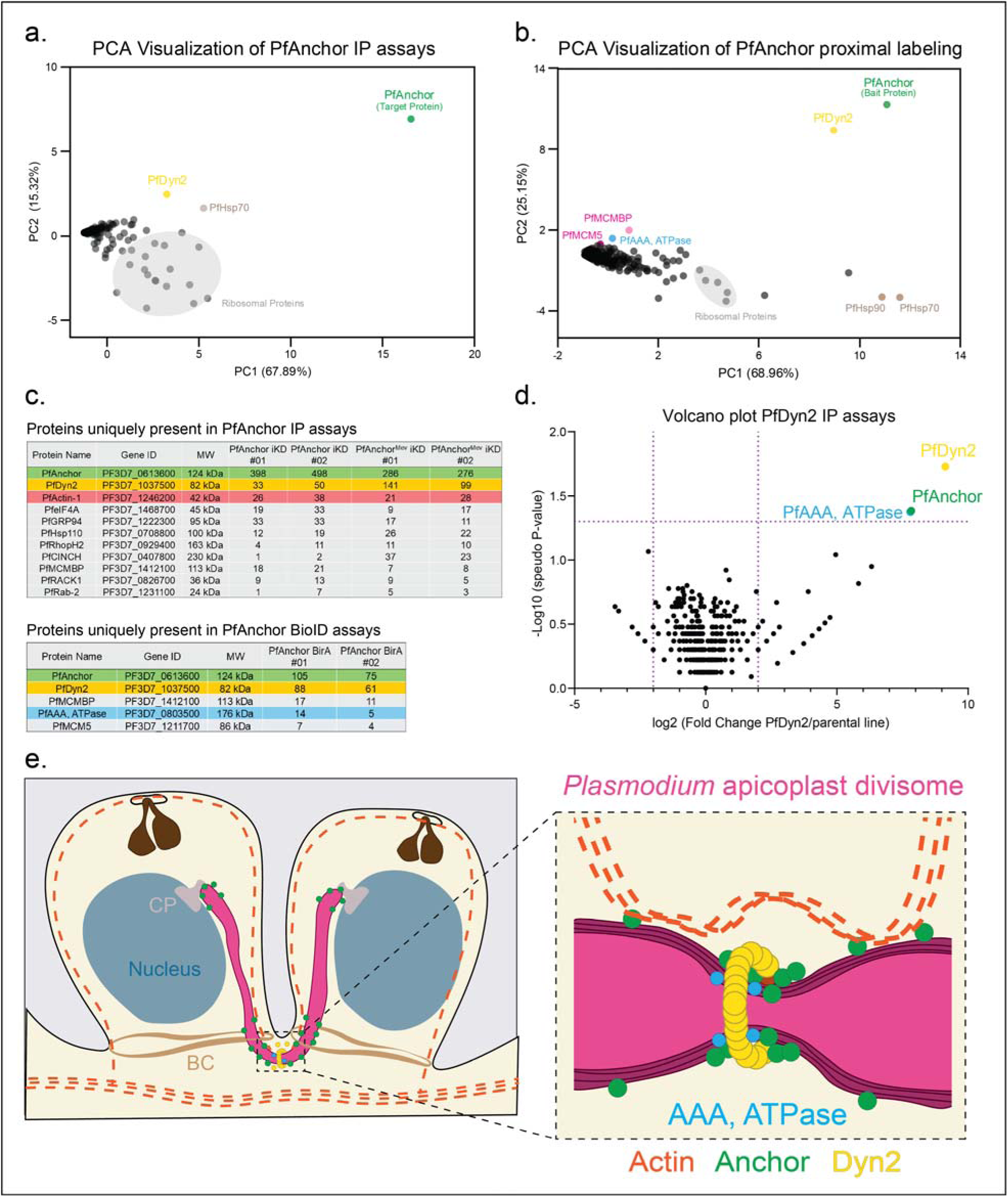
PfAnchor interacts with the dynamin-like GTPase PfDyn2, supporting a role in apicoplast division. **a.** Principal component analysis (PCA) of mass spectrometry data from PfAnchor immunoprecipitation (IP) assays, performed in biological duplicates using the PfAnchor iKD and PfAnchor^Mev^ iKD cell lines (as listed in c). PfDyn2 is identified as the primary interacting partner of PfAnchor, while PfHsp70 is also detected as a potential secondary interactor. Ribosomal proteins (gray) cluster separately as background. **b.** PCA of PfAnchor proximity labeling (BioID) mass spectrometry data. PfDyn2 is enriched in close proximity to PfAnchor, supporting its interaction. Additionally, components of the MCM complex are detected, though their functional relevance remains unclear. Tables listing proteins uniquely identified in PfAnchor IP and PfAnchor BioID (**c**) datasets compared to control samples. PfDyn2 is consistently enriched across both methods, further supporting its role as the primary interactor of PfAnchor. **d.** Volcano plot of mass spectrometry from PfDyn2 IP, performed in biological duplicates using the PfDyn2-3HAapt cell line. PfAnchor is identified as a primary interactor, along with PfAAA, ATPase, also identified in the PfAnchor proximal labeling experiments. **e**. Model of PfAnchor’s role in apicoplast division and inheritance. **Left:** During cytokinesis, the apicoplast (pink) is positioned adjacent to the centriolar plaque (CP) and basal complex (BC), where PfAnchor (green) localizes along the apicoplast membrane. PfDyn2 (yellow) is recruited to the fission site, potentially through PfAnchor, while actin (dashed orange lines) forms a dynamic network [61] that may contribute to membrane remodeling and organelle segregation. **Right:** Close-up view of apicoplast fission, where PfDyn2 assembles at the division site, driving membrane constriction. PfAnchor may function as a tethering factor, linking the fission machinery to the apicoplast membrane and ensuring proper inheritance of the organelle by daughter parasites.

To complement our IP approach and further define the PfAnchor interactome, we performed proximity-dependent biotin labeling (BioID), followed by mass spectrometry. BioID enables the identification of proteins in close spatial proximity to PfAnchor, including transient or membrane-associated interactors. Two independent biological replicates of BioID with PfAnchor-BirA-HA (**Figure S14**) were performed, with parental untagged parasites serving as a control to account for background labeling. PCA analysis of BioID mass spectrometry data demonstrated that PfAnchor clustered distinctly from background interactors, with PfDyn2 emerging as the most enriched protein in proximity to PfAnchor (**Figure 6b, Supp Data 3**). This further supports a functional association between PfAnchor and PfDyn2, consistent with the results of our immunoprecipitation experiments. Additional proteins detected in the BioID dataset included components of the MCM complex, though their potential relevance to apicoplast function remains unclear. PfDyn2 was consistently enriched in both BioID and IP datasets, reinforcing its role as PfAnchor’s primary interactor (**Figure 6c**). In addition, immunoprecipitation experiments using the parasite line NF54attB-PfDyn2-3HA^apt^ [27](PfDyn2-3HA^apt^) were performed to confirm association with PfAnchor. PCA of the mass spectrometry data confirmed PfAnchor as a primary interactor with PfDyn2 and identified a putative member of the AAA family ATPase that was also identified in the PfAnchor BioID (**Figure 6b-d**). This strong association suggests that PfAnchor may act as an adaptor or regulator in apicoplast fission, potentially recruiting PfDyn2 to the organelle membrane. Together, these findings support a model in which PfAnchor serves as a key adaptor protein, bridging cytoskeletal and membrane remodeling factors to facilitate apicoplast fission (**Figure 6e**).

## Discussion

This study reveals a fundamentally new mechanism of parasite death and identifies PfAnchor as an essential adaptor that enables apicoplast fission in *Plasmodium falciparum*. We show that PfAnchor localizes to the apicoplast throughout the intraerythrocytic cycle and is required for its division and inheritance. Conditional depletion of PfAnchor disrupts apicoplast fission, resulting in physically tethered merozoites that fail to complete cytokinesis and cannot invade new red blood cells. This phenotype demonstrates, for the first time, that failure to divide the apicoplast can directly impair cell division and lead to same-cycle parasite death. Additionally, we highlight the potential of targeting apicoplast morphology as a novel strategy for inducing same-cycle parasite death, positioning PfAnchor as a promising therapeutic target for malaria intervention.

### PfAnchor’s Structural Features and Functional Implications

Alphafold3 predictions and domain analysis suggest that PfAnchor contains a pleckstrin homology (PH) domain, a common motif for membrane binding. Given that the apicoplast membrane is enriched in PI3P [63, 64], it is plausible that PfAnchor’s PH domain mediates its localization. Additionally, PfAnchor’s N-terminal region contains a hydrophobic helix, which carries a mutation linked to resistance against actin-polymerization inhibitors [65]. The co-occurrence of mutations in PfAnchor and profilin (a key actin regulator) in parasites resistant to MMV020291, an antimalarial that targets actin polymerization, suggests a functional link between PfAnchor and actin dynamics [65]. Actin dynamics are increasingly recognized as important for apicoplast segregation, and our identification of PfActin1 as a PfAnchor interactor supports this idea [52, 61]. In other eukaryotic systems, profilin-actin interactions regulate fission events by remodeling membrane-associated cytoskeletal networks [66–68]. By analogy, PfAnchor may facilitate actin polymerization at the apicoplast fission site, potentially through indirect interactions with profilin. Future studies should investigate whether targeted disruption of PfAnchor’s hydrophobic helix affects actin-mediated apicoplast segregation.

### PfAnchor’s Role in Organelle Structure and Function

Despite its role in apicoplast fission, PfAnchor is not required for overall apicoplast biogenesis. Parasites lacking PfAnchor remain viable when the metabolic function of the apicoplast is bypassed using mevalonate supplementation, and our U-ExM data show that viable PfAnchor deficient parasites do not contain an apicoplast branching structure at day 8 post aTC removal (**Figure S11**). Furthermore, treatment with azithromycin, which eliminates the branched apicoplast structure, rescued the growth defects of PfAnchor-deficient parasites. These findings confirm that PfAnchor’s primary function is in organelle fission rather than metabolism.

Cryo-electron tomography of PfAnchor-deficient and control parasites further revealed the presence of distinct electron-dense structures within the apicoplast. We identified two main morphologies: “dense buds,” which are small, localized densities between membranes, and “enclosed buds,” which are larger, invaginated membrane profiles. Similar membranous profiles—sometimes described as tubular whorls—have been reported in apicoplasts of *Plasmodium* and *Toxoplasma*, though their functional role remains unresolved [56, 69, 70]. Comparative plastid systems, such as etioplasts, also contain analogous intra-organelle structures, including protein complexes, plastoglobuli, and tubular arrays involved in membrane remodeling and metabolic scaffolding [71]. The persistence of these features in the *Plasmodium* apicoplast, independent of PfAnchor expression, raises intriguing questions about their function in membrane remodeling, metabolite transport, or structural maintenance of the apicoplast’s four-membrane architecture. Whether these intramembrane structures persist within the apicoplast-like vesicles remains an open question. Further investigation will be required to determine whether these features are involved in apicoplast homeostasis, division, or are vestiges of its photosynthetic ancestry.

### Mitochondrion-Apicoplast Tethers and Organelle Inheritance

Our findings provide the first high-resolution evidence that mitochondrion-associated apicoplast-like vesicles are selectively inherited [72], reinforcing the functional dependency between these organelles [54, 56]. While most vesicles were excluded from daughter cells, the retained subset consistently colocalized with mitochondria (**Figures 5, S12, S13**). This strong spatial association suggests a role for physical tethering between the apicoplast and mitochondrion in ensuring faithful organelle inheritance. Moreover, the molecular components mediating this tethering are likely preserved on apicoplast-like vesicles that are inherited. Consistent with this idea, electron-dense structures at the interface between the mitochondrion and apicoplast have recently been described by cryo-ET in *P. falciparum*, suggesting the presence of defined tethering complexes at points of close membrane contact [56]. While we did not specifically image vesicles in contact with mitochondria by cryo-ET in this study, it remains an open and important question whether these dense tethering structures persist in the apicoplast-like vesicles that are inherited following AZ treatment. Interestingly, PfAnchor depletion resulted in a slight but significant increase in mitochondrion-associated vesicles (**Figure 5e**), indicating that while PfAnchor is not required for inheritance, its loss may alter the balance between vesicle retention and clearance. The nature of the mitochondrion–apicoplast tethers remains to be elucidated, but it is likely mediated by tethers that help coordinate spatial organization during division. Are these tethers simple structural linkages, or true membrane contact sites that enable metabolite exchange or signaling? The role of the cytoskeleton, particularly actin, in stabilizing these tethers warrants further investigation, as cytoskeletal elements are known to contribute to apicoplast segregation in *Plasmodium* and other apicomplexans[52, 61].

### PfAnchor Interacts with PfDyn2 and the Cytoskeleton

To elucidate the molecular role of PfAnchor, we combined immunoprecipitation and proximity labeling, which consistently identified PfDyn2 as its main interactor (**Figure 6**). Reciprocal immunoprecipitation using the PfDyn2-3HAapt line confirmed PfAnchor as a major PfDyn2 partner (**Figure 6d**). PfDyn2, a dynamin-related GTPase required for apicoplast and mitochondrial fission [27, 28], was enriched in both datasets, supporting a model where PfAnchor acts as a dynamin adaptor, recruiting PfDyn2 to the apicoplast membrane to facilitate fission. Notably, PfDyn2 immunoprecipitation also enriched a putative AAA-ATPase (Pf3D7_0803500) that appeared in the PfAnchor proximity labeling but not in its immunoprecipitation, indicating that this protein interacts directly with PfDyn2, not PfAnchor. Since Dynamin-AAA-ATPase interactions modulate mitochondrial fission in yeast [73], and PfDyn2 controls mitochondrial fission [27, 28], further research should investigate whether this AAA-ATPase contributes to an apicoplast or mitochondrial divisome in *Plasmodium*.

Additionally, we identified interactions between PfAnchor and cytoskeletal components, including PfActin1 and PfCINCH, suggesting a potential role in coordinating apicoplast fission with parasite cytoskeletal dynamics [61, 62]. Notably, the basal complex, which facilitates cytokinesis, may also coordinate organelle fission. Positioned at the parasite plasma membrane, it could serve as a platform for synchronizing apicoplast and mitochondrial division with daughter cell separation. Future studies should examine whether basal complex components, such as PfCINCH, actively remodel membranes or anchor fission machinery at the apicoplast and mitochondria during schizogony. Additionally, PfHsp70 (PF3D7_0818900), identified in our interactome analysis, could stabilize the fission complex, as its homologs in other eukaryotes support mitochondrial and chloroplast division by assisting in protein complex assembly and function [60, 73].

Finally, we identified PfAnchor via proximal labeling using PfMCMBP, a DNA helicase that functions during DNA replication [43]. While reciprocal proximity labeling with PfAnchor as bait confirmed the presence of PfMCMBP and PfMCM5, our immunoprecipitation experiments did not recover these proteins, suggesting that they are not direct binding partners of PfAnchor. Instead, their detection in proximity labeling assays likely reflects their spatial localization within the parasite during schizogony rather than a functional interaction. The apicoplast and PfAnchor are positioned near the centriolar plaque (CP) immediately before segmentation (**Figure S6**), and the CP is surrounded by nuclear pore complexes, indicating active nucleocytoplasmic transport in this region [74]. Given this, it is plausible that the MCM complex, which shuttles between the cytoplasm and nucleus during DNA replication, transiently occupies a similar cellular space. While a direct functional relationship between PfAnchor and the MCM complex remains unlikely, these findings highlight the complex organization of cellular structures during *P. falciparum* blood-stage parasites replication and underscore the utility of proximity labeling in capturing transiently colocalized proteins within the parasite.

### PfAnchor in Other Developmental Stages

Although our study establishes PfAnchor’s role in blood-stage parasites, its expression in other life cycle stages suggests broader functions. Transcriptomic data indicate that PfAnchor is expressed during gametocytogenesis, where male and female gametocytes harbor a single, non-branching apicoplast [75, 76]. Whether PfAnchor plays a role in gametocyte development or is dispensable at this stage remains unclear. By contrast, during mosquito and liver stages, parasites undergo extensive replication, and their apicoplasts adopt a branching tubular morphology similar to blood stages [77]. This implies that apicoplast fission must occur to ensure organelle inheritance during liver merozoite and sporozoite formation. Investigating PfAnchor’s function in these stages could reveal conserved mechanisms of apicoplast division throughout the parasite life cycle and identify new transmission-blocking targets.

### Conservation and Lineage-Specificity of the Apicoplast Divisome

To place our findings in an evolutionary context, it is important to ask whether the apicoplast divisome is conserved across apicomplexans or represents a *Plasmodium*-specific innovation. Like other apicomplexans that lack the bacterial FtsZ/ARC5 division systems, *Toxoplasma gondii* relies on the dynamin-related protein DrpA for apicoplast fission, with no adaptor protein identified to date [78]. In contrast, *P. falciparum* uses PfDyn2 for mitochondrial and apicoplast fission [27]. Comparative insights from lineages that have retained their apicoplasts, such as *Toxoplasma*, and from those that have secondarily lost them, such as *Cryptosporidium* [79], offers a unique opportunity to distinguish conserved from lineage-specific mechanisms of plastid division.

Our BLASTp survey across representative apicomplexan genomes did not find clear homologs of PfAnchor or the putative apicoplast AAA ATPase outside plastid-bearing lineages (**Figure S16**). This pattern suggests that these proteins are specialized for apicoplast division in *Plasmodium*. However, it also highlights the limitations of sequence-based inference, as highly diverged adaptors may go undetected. Future experiments, such as immunoprecipitation of TgDrpA in *T. gondii* to identify its interacting partners, could help uncover functional analogs of Anchor or AAA ATPase in other apicomplexans and provide a more complete picture of the plastid fission machinery.

Taken together, our findings identify PfAnchor as a lineage-specific adaptor that facilitates organelle-specific recruitment of PfDyn2 to the apicoplast, while mitochondrial fission remains under the control of yet-to-be-identified factors. The specific requirement for PfAnchor likely reflects the complex four-membrane structure of the apicoplast, which originates from a red algal endosymbiont, and highlights how conserved dynamin-related proteins can be paired with different adaptors to reshape organelles of diverse evolutionary origins. Comparisons with photosynthetic relatives like dinoflagellates emphasize how the same dynamin machinery has been adapted to serve plastids with different functions, from metabolism-centric apicoplasts in parasites to photosynthetic organelles in free-living species. Since apicoplast division is essential for parasite survival and has no equivalent in human cells, targeting PfAnchor or its associated machinery presents a promising new approach for antimalarial therapy.

## Materials and Methods

### Parasite culture

The 3D7 *Plasmodium falciparum* strain, obtained from the Walter and Eliza Hall Institute (Melbourne, Australia) was cultured in human O^+^ RBCs at 4% hematocrit in RPMI-1640 containing 25 mM HEPES, 50 mg/L hypoxanthine, 0.21% sodium bicarbonate, and 0.5% w/v Albumax II [80]. Cultures were incubated at 37 °C while shaking in a gas mixture of 1% O_2_, 5% CO_2_, and 94% N_2_ as previously described The PfAnchor iKD (smV5-Tet Pf3D7_0613600) and PfAnchor^Mev^ iKD (smV5-Tet Pf3D7_0613600 in PfMev background) cell lines were maintained under selection of 2.5 nM WR99210 (5 nM, Jacobs Pharmaceutical) and supplemented with 500 nM anhydrotetracycline (aTC). PfDyn2-3HAapt was maintained under selection of blasticidin (2.5 µg/mL, InvivoGen) and supplemented with 500 nM anhydrotetracycline (aTC).

### Plasmid generation and transfection

To generate the PfMCMBP-BirA-HA cell line for proximal labeling, the homology region of pSAB60 (PfMCMBP-3HADD [43]) was cut with Not1/Xho1 and cloned into our BirA plasmid (pRR28) [81] to create pSAB99. To generate the PfMCMBP-BirA-HA strain, 100 µg of pSAB99 plasmid was transfected into synchronized ring staged parasites using electroporation. Upon transfection, stable single crossover parasites were selected by cycling 2.5 nM WR99210 on and off as previously described [82]. Integration of pSAB99 plasmid into the genome of 3D7 was confirmed via PCR using primer pairs oSAB504/oJDD44, oSAB505/oJDD44, and oSAB504/oSAB506 **(Figure S1**).

To create the Pf3D7_0613600-smV5-Tet (PfAnchor iKD) plasmid, the Pf3D7_0613600 5’ and 3’ homology regions were PCR amplified from 3D7 genomic DNA with oligonucleotides oJDD4308/4310 and oJDD4307/4309 respectively. The two pieces were fused together using Sequence Overlap Extension PCR (SOE-PCR) using oJDD4803/4309 and the piece was digested using Not1/Nco1 and ligated onto pRR92 [62] (which contains the smV5 epitope tag, ten copies of the Tet-aptamer for the TetR-DOZI aptamer knockdown system, and the expression cassette for human dihydrofolate reductase (hDHFR)) to form pSAB177. To create Pf3D7_0613600 targeting guide RNA plasmid, oJDD4347/4348 were annealed, phosphorylated, and ligated into BpiI-digested pRR216 [27] to generate pSAB175. All oligonucleotide sequences are shown in **Table S1**. To generate the PfAnchor iKD strain, 100 µg of pSAB177 plasmid was linearized with Stu1 and transfected into 3D7-Cas9 parasites, along with 100 µg of pSAB175 and supplemented with 500 nM aTC. 24 hours following transfection parasites were treated with 5 nM WR99210 for 7 days before maintaining on 2.5 nM WR99210 and 500 nM aTC until resistant parasites were detected. Integration of pSAB177 plasmid into the genome of 3D7 was confirmed via PCR using primer pairs oSAB243/oJDD2933, oSAB367/oSAB2933, oSAB243/oSAB366, and oSAB367/oSAB366. Primer binding locations and predicted PCR product sizes are listed in **Figure S4a-b**.

To generate the PfAnchor^Mev^ iKD strain, 350 μL of RBCs were electroporated using the GenePulser XCell system (Bio-Rad) with 75 μg each of pSAB175 and linearized pSAB177 obtained by digestion with EcoRV. The electroporated RBCs were combined with 1 mL of PfMev^attB^ parasite culture at 3% parasitemia and cultured in 10 mL of CMA (Complete Medium with Albumax) for two days. Subsequently, the cultures were transferred to selective medium containing 2.5 nM WR99210 for seven days. Afterward, the cultures were returned to CMA until parasites were detected on blood smears, at which point WR99210 was reintroduced to the medium. The transgenic parasite culture was cloned by limiting dilution. Clones were screened for an intact aptamer array via PCR amplification using primers oSAB249 and oJDD44. Clone F3 was selected for further experiments (**Figure S15**). Integration of pSAB177 plasmid into the genome of NF54-PfMev parasites [50] was confirmed via PCR using primer pairs oSAB243/oJDD2933, oSAB367/oSAB2933, oSAB243/oSAB366, and oSAB367/oSAB366. Primer binding locations and predicted PCR product size found in **Figure S4a, c**.

To generate the PfAnchor-BirA-HA proximal labeling plasmid, the homology region from pSAB177 was amplified using oSAB356/358 and the piece was digested using Not1/Xho1 and ligated into pSAB99 to create pSAB233. To generate the PfAnchor-BirA-HA strain, 100 µg of pSAB233 plasmid was linearized with Stu1 and transfected into 3D7-Cas9, along with 100 µg of pSAB175. 24 hours following transfection parasites were treated with 5 nM WR99210 for 7 days before maintaining 2.5 nM WR99210 until resistant parasites were detected. Integration of pSAB233 plasmid into the genome of 3D7 was confirmed via PCR using primer pairs oSAB243/oJDD44, oSAB243/oSAB366, oSAB367/oJDD44, and oSAB367/oSAB366. Primer binding locations and predicted PCR product size are found in **Figure S14**. All parasite lines used in this study are listed in **Table S2**.

### Growth analysis

For parasite growth analysis, parasite cultures were synchronized at the schizont stage by either density centrifugation using 60% Percoll PLUS or by magnetic separation using MACs columns, incubated at 37 °C for 2-3 hours with fresh erythrocytes, and then newly invaded ring stage parasites were purified by treatment with 5% w/v sorbitol. In three biological replicates, cultures were plated and incubated at 37 °C at 4% hematocrit in the presence or absence of aTC. Over the course of 96 hours (2 intraerythrocytic cycles) timepoints were taken at 0, 48, and 96 hours and the parasitemia was determined by counting a minimum of 400 RBCs.

### Sodium carbonate extraction

For the sodium carbonate extraction assay, the protocol from Liffner *et al* was used [83]. Saponin-lysed parasite pellets from 10 mL of high parasitemia late schizonts (40-48 hpi) were resuspended in 100 µL of Milli-Q water and snap-frozen using liquid nitrogen four times before passing through a 28 gauge needle. Samples were centrifuged for 10 minutes at 18,000 x g at 4 °C, with the supernatant reserved as the hypotonic (water soluble) sample. The pellet was washed twice with 500 µL Milli-Q water and once with 500 µL 1x PBS before the pellet was resuspended in 100 µL 0.1 M sodium carbonate (Na_2_CO_3_) and incubated in ice for 30 minutes. The sample was centrifuged at 18,000 x g at 4 °C for 5 minutes and the supernatant was reserved as the sodium carbonate (peripheral membrane) sample. The remaining pellet was washed 3 times in 100 µL 1x PBS before resuspending the pellet in 100 µL ice cold 0.1% Triton X-100/PBS and incubated for 30 minutes on ice. Samples were centrifuged at 18,000 x g at 4 °C and the supernatant was reserved before the pellet was washed and resuspended in 100 µL 1x PBS. Samples were analyzed by western blotting.

### Proteinase K protection

For Proteinase K protection assay, the protocol from Liffner *et al* was used [83]. 3x 10 mL cultures of schizonts (36-48 hpi) were lysed with 0.15% saponin/PBS-protease inhibitors (Sigma S8820-20 TAB, 1 tab for 100 mL PBS) for 10 minutes before centrifuging at 18,000 x g for 10 minutes at 4 °C. Pellets were washed 3x in 1x PBS-protease inhibitors before further treatment. Three treatments were performed: one tube treated with 250 µL SOTE buffer (0.6 M Sorbitol, 20 mM Tris HCl, 2 mM EDTA) alone. A second tube was treated with 250 µL 0.02% w/v digitonin in SOTE for 10 minutes on ice before centrifuging at 800 x g for 10 minutes at 4 °C. The resulting pellet was washed once with SOTE buffer. A third tube of sample was treated with digitonin followed by treatment with 0.1 mg/mL Proteinase K in SOTE for 30 minutes on ice. The sample was then centrifuged at 18,000 x g for 10 minutes at 4 °C before inactivating the Proteinase K by treatment of the pellet with 100 µL of 5 mM PMSF (ThermoFisher #36978) in SOTE buffer for 10 minutes on ice. Samples were analyzed by western blotting.

### Western Blot

Protein samples were collected by saponin lysis with 0.15% w/v saponin/PBS-protease inhibitors for 10 minutes on ice, parasite material was pelleted by centrifugation before washing 3x with PBS-protease inhibitors. Parasite pellets were resuspended in Laemmli sample buffer + 5% v/v β-mercaptoethanol (Aldrich), heated at 37 °C for 1 hour with vortexing every 30 minutes, and separated by size using BioRad 4-20% Mini-PROTEAN TGX stain free gels (BioRad Cat. #4568095) at 200 V for 40 minutes. Proteins were transferred to a nitrocellulose membrane using a BioRad Transblot Turbo Transfer System with the MixedMW setting (25 V, 1.3 A, 7 minutes) before blocking using 1:5 BioRad EveryBlot Blocking Buffer (BioRad Cat. #12010020): PBS for 30 minutes at room temperature. Membranes were incubated in primary antibodies (listed in Table S3) overnight at 4 °C, washed 3x in 1x TBS+ 0.01% Tween20 before incubating with secondary antibodies (listed in Table S3) at room temperature for 45 minutes. Western Blots were visualized using a BioRad ChemiDocMP Imaging system. Western blot quantification was done using BioRad Image Lab V6.1 software.

### Live microscopy

Glass bottomed microscopy dishes (Cellvis D35-20-1.5-N) were coated with 0.5 mg/mL concanavalin A (Sigma, C0412-5MG) solution for 1 hour at 37 °C before 3x washes with Milli-Q water. Synchronous late stage +aTC or -aTC parasites were purified by MACs columns and treated with compound 1 [51] before being applied to the glass bottomed dishes and incubated for 3 hours at 37 °C to allow parasites to adhere. After 3 hours samples were washed with 1 mL complete phenol-red-free RPMI. Brightfield and fluorescent images were taken every 0.5-4.5 seconds for 30 minutes per sample using a Leica DMI6000 B microscope and a 63x objective with a numerical aperture of 1.4.

### Ultrastructure Expansion Microscopy (U-ExM)

12 mm round coverslips (Fisher, Cat# NC1129240) were treated with poly-D-lysine for 1 hour at 37 °C, washed twice with MilliQ water, and placed in the wells of a 24 well plate. Five hundred uL of parasite cultures adjusted to roughly 1% hematocrit was added to the wells containing a coverslip. Samples were allowed to settle for 30 minutes at 37 °C before culture supernatant was removed and 500 µL of 4% v/v PFA in 1x PBS was gently added along the side of the well and incubated at 37 °C for 20 minutes. Coverslips were washed once with 1x PBS before being treated with 500 µL of 1.4% v/v formaldehyde/2% v/v acrylamide (FA/AA) in PBS overnight at 37 °C. Gelation, denaturation, staining, and expansion of gels were performed as previously described [25]. Stained gels were imaged using a Zeiss LSM900 AxioObserver with an Airyscan 2 detector. Images were taken using a 63x Plan-Apochromat objective lens with a numerical aperture of 1.4.  All antibodies and their dilutions used in this study are listed in **Table S3**.

### Cryo-Electron Tomography

#### Sample preparation

PfAnchor iKD parasites were grown and synchronized as described above. Late schizont pellets were obtained via percoll purification and allowed to incubate in 5 mL of media with 2.5 μM Compound 1 [51] for 3 hours. Parasites were chemically fixed with 4% PFA/0.01% glutaraldehyde for 20 minutes before resuspending in PBS prior to cryo-electron tomography.

#### Cell vitrification

Quantifoil Cu R2/1 200 mesh grids were glow-discharged for 1 minute at 25 mA using an EmiTech K100X glow discharger (EMS). Fixed PfAnchor iKD parasites, prepared as described above, with or without aTC and arrested with Compound 1, were applied in 4 µL volumes to each grid for vitrification, then manually back-blotted on one side (at 22°C and 100% humidity) for 3-4 seconds, and plunge-frozen in a 1:1 mixture of liquid ethane and propane using a Vitrobot Mk IV (Thermo Fisher Scientific - TFS, Waltham, MA, USA). Plunge-frozen grids were clipped with auto-rings and stored in liquid nitrogen for loading.

#### Cryo-FIB Milling

Cryo-FIB milling was performed using an Aquilos 2, a cryo-dedicated DualBeam microscope (TFS). Cryo-grids were sputter coated with platinum for 10 seconds, followed by a ∼500 nm thick layer of platinum deposited by the gas injection system for 15 seconds, followed by another 10 seconds of platinum sputter coating. Cell clusters were milled to generate lamella at ∼150-200 nm thickness, using 0.1-0.3 nA ion beam current for rough milling and 30-50 pA for final polishing.

#### Cryo-ET

Cryo-FIB milled lamella grids were transferred to a Titan Krios 3Gi microscope (TFS), equipped with a 300 kV field emission gun, a Selectris energy filter and Falcon 4i direct electron detector. Tilt series were collected using Tomo5 software (TFS) at 42,000x magnification (2.9 Å pixel size) and the energy filter at 20.0 eV slit width. Tilt ranges were adjusted for each acquisition site to the minimum and maximum tilt angles between -60° and +60° that encompassed the feature of interest. Tilt series were acquired in 3° increments with each tilt image recorded over 5 movie frames at defocus values between -4 to -8 μm. The total dose applied for each tilt series was ∼100 e-/Å^2^.

#### Tomogram Reconstruction and Analysis

Movie frames for each tilt series were motion corrected using MotionCor2 [84] through Relion 3.1 [85] using 5x5 patches. Motion-corrected tilt series were generated using EMAN2. Tilt series alignment, tomogram reconstruction, and CTF estimation were performed using the automated EMAN2 pipeline [86] or IMOD [87, 88]. Feature segmentation was performed using neural network training in EMAN2 or manually using IMOD 3dmod. Tomogram features were visualized and analyzed using ChimeraX [89, 90].

### Mevalonate bypass experiments

#### Azithromycin treatment

To generate PfAnchor^Mev^ iKD parasites with disrupted apicoplasts, Clone F3 parasites were treated with 100 nM azithromycin (1× IC₅₀) (Sigma PZ0007) for one week, with continuous supplementation of 50 μM mevalonate (Sigma M4667). The untreated parental Anchor knockdown line served as a positive control for apicoplast genome detection (**Table S1**).

#### Measurement of parasite growth

To assess the dependence of apicoplast-intact and apicoplast-disrupted PfAnchor^Mev^ iKD parasites on mevalonate for survival, asynchronous parasites were washed three times with 10 mL of CMA to remove residual mevalonate and aTC. They were seeded in a 96-well plate at an initial parasitemia of 0.5% and a hematocrit of 2%, with a total volume of 200 μL per well. The parasites were then cultured under four conditions, with each condition tested in quadruplicate: (1) CMA with 50 μM mevalonate and aTC, (2) CMA with 50 μM mevalonate, (3) CMA with aTC, and (4) CMA alone. Cultures were incubated at 37°C in chambers gassed with 94% N₂, 3% O₂, and 3% CO₂. Every other day, 10 μL of each sample was collected, diluted 1:10 in phosphate-buffered saline (PBS), and stored in a 96-well plate at 4°C. To prevent overgrowth, cultures were simultaneously diluted at 1:8.

For growth curve determination, parasite samples were analyzed by flow cytometry over two intervals: days 0 to 4 and days 5 to 8. On day 4, 10 μL of each diluted sample was transferred to a new 96-well plate containing 100 μL of 1× SYBR Green (Invitrogen) in PBS per well. The plates were incubated at room temperature in the dark with gentle shaking for 30 minutes. After staining, 150 μL of PBS was added to each well to dilute excess dye. The samples were analyzed using an Attune Nxt flow cytometer (Thermo Fisher Scientific) with a 50-μL acquisition volume, a flow rate of 25 μL/min, and a total collection of 10,000 events per sample. The same protocol was repeated on day 8 for the second set of samples collected on days 6 and 8.

#### Analysis of apicoplast genome

To assess the integrity of the apicoplast genome after treatment with azithromycin, PfAnchor^Mev^ iKD parasites were grown for 5 days in the presence of 100 nM azithromycin and 50 µM mevalonate before aTC washout. Genomic DNA was extracted using QIAmp DNA blood mini kit (Qiagen, #51106) every two days up to day eight. Control samples from PfAnchor^Mev^ iKD not treated with azithromycin were also collected. PCR analysis was performed using primers oSAB484/oSAB485 (GAPDH, nuclear DNA), oSAB486/oSAB487 (tufa, apicoplast DNA), and oSAB488/oSAB489 (cytb3, mitochondrion DNA) as previously described [91].

### Immunoprecipitation and proximal labeling using PfAnchor as bait

#### Sample preparation

Protein extraction for both immunoprecipitation (IP) and BioID experiments was performed using the same lysis procedure. Parasites were cultured at 2% hematocrit and 4–5% parasitemia before harvesting. Red blood cells were lysed using 0.05% saponin in PBS supplemented with protease inhibitors (SigmaFast, Sigma S8820-20TAB) and incubated on ice for 10 minutes. Parasites were washed three times with ice-cold PBS + protease inhibitors by centrifugation at 10,000 g for 5 minutes at 4°C. The resulting pellets were resuspended in 1 mL PBS + protease inhibitors, centrifuged at 10,000 rpm for 2 minutes at 4 °C, and stored at -80 °C or processed immediately. Parasite pellets were lysed in 1 mL RIPA buffer (50 mM Tris-HCl pH 7.5, 150 mM NaCl, 1% NP-40, 0.5% sodium deoxycholate, 0.1% SDS) supplemented with protease inhibitors. Samples were incubated on a rotator at 4 °C for 30 minutes. To ensure efficient lysis, three rounds of sonication (30 seconds at 20 amplitude) were performed on ice, with 3-minute rest intervals between pulses. Lysates were cleared by centrifugation at 15,000 g for 30 minutes at 4 °C, and the supernatant was transferred to a fresh tube. For the MCMBP-BioID experiments, soluble and insoluble fractions were processed separately using the same protocol described in our previous study [43]. For immunoprecipitation, 50 µL of anti-V5 magnetic beads (Thermo, #88816) per sample were washed twice with 1 mL RIPA buffer and incubated with the cleared lysates overnight at 4 °C with constant rotation. Beads were collected using a magnetic stand, washed three times with 50 mM ammonium bicarbonate buffer, and resuspended for on-bead digestion and mass spectrometry analysis. For BioID, 50 µL of streptavidin magnetic beads (Thermo, #88816) were used instead of V5 magnetic beads. After overnight incubation, beads were washed following the same procedure as in IP before processing for mass spectrometry analysis.

#### On bead digests

After washing, beads were covered with 8 M Urea, 100 mM Tris hydrochloride, pH 8.5, reduced with 5 mM tris (2-carboxyethyl) phosphine hydrochloride (TCEP, Sigma-Aldrich Cat No: C4706) for 30 minutes at room temperature to reduce the disulfide bonds. The resulting free cysteine thiols were alkylated using 10 mM choloracetamide (CAA, Sigma Aldrich Cat No: C0267) for 30 minutes at RT, protected from light. Samples were diluted to 2 M Urea with 50 mM Tris pH 8.5 and proteolytic digestion was carried out with Trypsin/LysC Gold (0.4 µg, Mass Spectrometry grade, Promega Corporation Cat No: V5072) overnight at 35 °C. After digestion, samples were quenched with 0.4% trifluoroacetic acid (v/v, Fluka Cat No: 91699). Samples were either injected onto a trap column followed by analytical column or first subjected to a solid phase extraction clean up on Pierce C18 spin columns (Cat No 87870).

### Immunoprecipitation using PfDyn2 as bait

Two parasite lines, PfDyn2-3HA^apt^ and the parental wildtype NF54attB, were cultured and tightly synchronized to harvest late-stage schizonts. On the day of harvest, the cultures were collected and treated with saponin (0.05% Saponin/PBS) for 5 min. The parasite pellets were thoroughly washed with PBS to remove any residual hemoglobin. The parasite pellets (∼ 100 µL of each line) were then solubilized in four volumes of the buffer containing 1% glycol-diosgenin (GDN), 50 mM NaCl, 10 mM MgCl_2_, 200 mM 6-aminocapronic acid, 50 mM Imidazole/HCl, 10% Glycerol and 1x Protease inhibitor cocktail. Post overnight solubilization on a rotator at 4 °C, the samples were spun down at 14,000 rpm at 4°C for 20 min to collect the supernatants. The protein concentrations of the supernatants were determined by the Pierce BCA Protein Assay (catalog, A55864). Meanwhile, Pierce anti-HA coated Magnetic beads (catalog, 88837; 50 µL for each parasite line) were washed three times with PBS using the DynaMag™-2 magnet (catalog, 12321D). The beads were then equilibrated with 500 µL of solubilization buffer without GDN for a few min. After removal of the equilibration buffer, the beads were loaded with supernatants of PfDyn2-3HA^apt^ or NF54attB (1 mg total protein for each line). The samples were rotated at 4 °C for overnight then washed three times with PBS using the DynaMag™-2 magnet. The washed beads were resuspended in a small volume of PBS (50 µL) and shipped on ice packs to the Proteomics & Mass Spectrometry Core Facility at University of South Florida. The samples were further digested with trypsin and analyzed by Mass Spectrometry following standard protocols.

#### Liquid Chromatography Mass Spectrometry

Approximately 1/15^th^ of each IP sample was loaded onto a 5 cm C18 trap column Acclaim™ PepMap™ 100 (3 µm particle size, 75 µm diameter; Thermo Scientific, Cat No: 164946) followed by a 25 cm EASY-Spray column (Thermo Scientific, Cat No: ES902) and analyzed using a Q-Exactive Plus mass spectrometer (Thermo Fisher Scientific) operated in positive ion mode. Solvent B was increased from 5%-35% over 100 min, to 90% over 2 min, back to 3% over 2 minutes (Solvent A: 95% water, 5% acetonitrile, 0.1% formic acid; Solvent B: 100% acetonitrile, 0.1% formic acid). A data dependent top 15 method was used with MS scan range of 350-2000 m/z, resolution of 70,000, automatic gain control (AGC) target 3e6, maximum injection time (IT) of 200 ms. MS2 resolution of 17,500, scan range of 200-2000 m/z, normalized collision energy of 30, isolation window of 4 m/z, target AGC of 1e5, and maximum IT of 150 ms. Dynamic exclusion of 10 sec, charge exclusion of 1, 7, 8, >8 and isotopic exclusion parameters were used. For samples post C18 cleanup, after drying in a speed vacuum, samples were resuspended in 25 µL 0.1 % FA. Approximately 1/5^th^ of each sample was then injected using an EasyNano1200 LC coupled to an Exploris 480 orbitrap mass spectrometer (Thermo Fisher Scientific)). Solvent B (80% Acetonitrile, 0.1 % FA) was increased from 8-35 % over 90 minutes, increased from 35-65 % over 15 min, increased to 85% over 5 min, held at 85% for 5 min, and decreased to 4% over 5 min. The mass spectrometer was operated in positive ion mode, advanced peak determination on, default charge state of 2 and user defined lock mass of 445.12003. 4 second cycle time was used with MS1 parameters of scan range 375-1500 m/z: orbitrap resolution of 60,000, standard AGC, automatic max IT, and RF lens of 40%. Monoisotopic peak determination was set to peptide with a minimum intensity filter of 5e3, charge state filter of 2-7, and dynamic exclusion of 30 s with 5 ppm mass tolerance. MS2 parameters included an isolation window of 4 m/z, normalized high energy dissociation energy of 30 %, orbitrap resolution of 15,000, user defined first mass of 110 m/z, standard AGC target and auto max IT. Data were recorded using Tune application 4.2.362.42.

#### PfDyn2 Data Analysis

A Volcano Plot was generated from the mass spectrometry dataset. For each protein, the average number of unique peptide in the PfDyn2 immunoprecipitations was divided by the average unique peptide count in NF54attB controls, and the ratio was Log_2_ transformed (X-axis). Positive values indicate enrichment in PfDyn2 samples, negative values indicate enrichment in controls, and values near zero indicate no enrichment. The Y-Axis represents the confidence of enrichment, calculated as 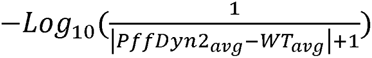. Larger values reflect higher confidence, while lower values correspond to greater uncertainty.

### Homology search

We searched VEuPathDB (BLASTP, default settings) against the reference proteomes shown in **Fig. S16** using three *P. falciparum* queries: Dyn2 (PF3D7_1037500), AAA+ ATPase (PF3D7_0803500), and Anchor (PF3D7_0613600). Candidates were retained if they met: (i) E-value ≤ 1e-50 (relaxed to 1e-40 for Anchor due to rapid divergence), (ii) alignment coverage ≥ 60%, and (iii) reciprocal best hit back to the *P. falciparum* query. For the AAA+ ATPase, we additionally required matching domain architecture and excluded other AAA+ families by annotation/domain (InterPro): FtsH/AFG3 (M41 peptidase), proteasome Rpt, CDC48/p97, VPS4 (MIT), and DEAD/DEAH helicases. Presence or absence per species was determined from the highest-scoring RBH-supported hit that met these filters. Under these criteria, Dyn2 homologs were found to be broadly distributed across apicomplexans; however, Anchor and the queried AAA+ ATPase were retained only in *Plasmodium*.

#### Data analysis

Data were analyzed using Proteome Discoverer 2.5.0.400 (Thermo Fisher Scientific). *Plasmodium falciparum* 3D7 and *Homo sapiens* reference proteome databases (downloaded from Uniprot on 04/06/2021 with 5381 sequences and on 05/13/2022 with 78806 sequences respectively), plus common laboratory contaminants (73 sequences) were searched using SEQUEST HT. Precursor mass tolerance was set to 10 ppm and fragment mass tolerance set at 0.02 Da with a maximum of 3 missed cleavages. A maximum of 3 modifications were allowed per peptide. Dynamic modifications included methionine oxidation; phosphorylation on serine, threonine, and tyrosine; dynamic protein terminus modifications were acetylation, met-loss, and met-loss plus acetylation. Static modifications were carbamidomethylation on cysteines. Percolator false discovery rate (FDR) filtration of 1% was applied to both the peptide-spectrum match and protein levels. Search results were loaded into Scaffold Q + S Software (version 5.2.2, Proteome Software, Inc) for visualization.

## Data availability

Raw and processed mass spectrometry data have been uploaded to ProteomeXchange with accession PXD062126. Dyn2 IP mass spectrometry raw and processed data have been uploaded to MassIVE with accession MSV000099148.

## Author contributions

**James Blauwkamp:** Methodology, Formal analysis, Investigation, Data curation, Writing - Original Draft, Writing - Review & Editing, Visualization.

**Krithika Rajaram:** Methodology, Investigation, Data curation, Visualization, Review & Editing

**Sophia R. Staggers:** Methodology, Investigation, Data curation, Visualization, Review & Editing.

**Oliver Harrigan:** Methodology, Data curation.

**Emma H. Doud:** Methodology, Formal analysis Data curation, Review & Editing

**Wei Xu:** Methodology, Investigation, Data curation

**Hangjun Ke:** Methodology, Investigation, Data curation, Review & Editing

**Sean T. Prigge**: Review & Editing, Funding Acquisition.

**Stella Y. Sun:** Methodology, Formal analysis, Investigation, Data curation, Resources, Review & Editing, Supervision.

**Sabrina Absalon:** Conceptualization, Methodology, Formal analysis, Investigation, Data curation, Visualization, Resources, Writing – Review & Editing, Supervision, Project administration, Funding Acquisition.

## Acknowledgements

We thank Michael Holmes and Anat Florentin for insightful discussions and review of the manuscript. We thank BEI Resources (NIAID, NIH), the European Malaria Reagent Repository, Vasant Muralidharan and Arnab Pain for providing the antibodies. We are grateful to A.J. Baucum for generously allowing us access to his equipment for western blot analyses. We also thank Jeffrey D. Dvorin for creating a stimulating and supportive research environment during Sabrina Absalon’s postdoctoral training, which encouraged her scientific independence, enabled her to develop side projects such as Anchor, and allowed her to bring these datasets to her independent laboratory. Mentors like him are critical to supporting the success and independence of early-career scientists. This research utilized PlasmoDB, a VEuPathDB resource [92]. This research was supported by Grant No. 2023198 from the United States-Israel Binational Science Foundation (BSF) and was made possible by an award from the Indiana University School of Medicine (S.A., BRG award no. EPAR2216). The content is solely the responsibility of the authors and does not necessarily represent the official views of the Indiana University School of Medicine. This research was also supported by S.T.P. from the National Institutes of Health (R01 AI125534), the Johns Hopkins Malaria Research Institute, and Bloomberg Philanthropies, with S.T.P. as principal investigator. The mass spectrometry analyses for PfAnchor experiments were performed by the Indiana University School of Medicine Proteomics Core, supported by start-up funds from S.A. We also gratefully acknowledge Dr. Dale Chaput at the University of South Florida for his assistance with protein mass spectrometry of the PfDyn2 immunoprecipitation samples. Acquisition of the IUSM Proteomics core instrumentation used for this project was provided by the Indiana University Precision Health Initiative. The proteomics work was supported, in part, by the Indiana Clinical and Translational Sciences Institute (funded in part by Award Number UL1TR002529 from the National Institutes of Health, National Center for Advancing Translational Sciences, Clinical and Translational Sciences Award) and, in part, by the IU Simon Comprehensive Cancer Center Support Grant (Award Number P30CA082709 from the National Cancer Institute). The Pittsburgh Center for CryoEM (RRID:SCR_025216) used for data collection in this project was supported, in part, by the University of Pittsburgh, the School of Medicine, the Department of Structural Biology, and the National Institutes of Health (grants S10-OD-019995 and S10-OD-025009).

