## Supplementary material for "An Essential Adaptor for Apicoplast Fission and Inheritance in Malaria Parasites": PfAnchor Supp Figures Resub.docx

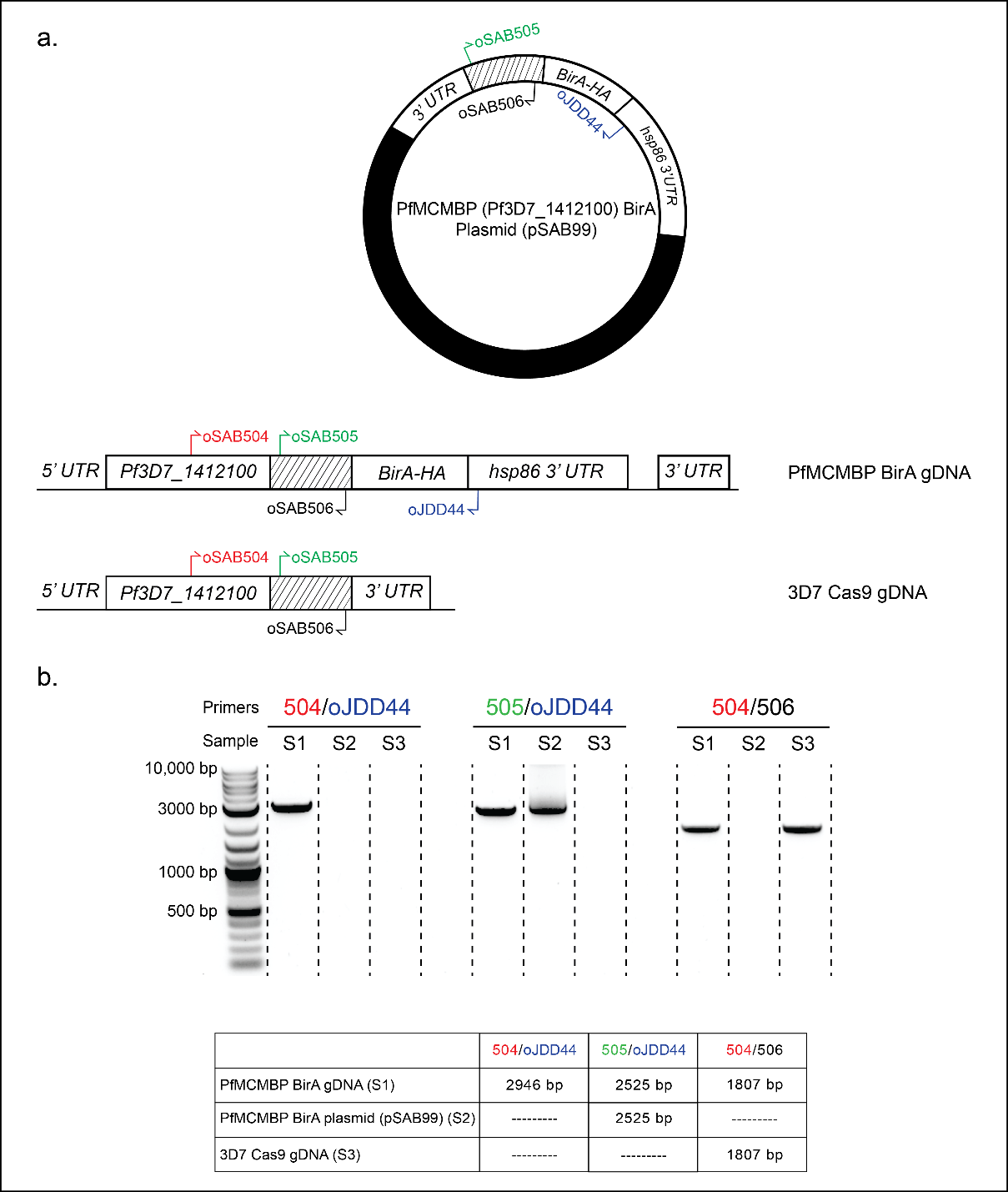

**Figure S1: Generation and confirmation of the PfMCMBP BirA-HA cell line.**

**a.** Schematic representation of the PfMCMBP (Pf3D7_1412100) plasmid and locus before and after integration of the BirA-HA tag. 3D7 Cas9 parasites were transfected with pSAB99 containing a 3’ homology region (marked with hashed lines) of PfAnchor, followed by a C-terminal BirA-HA. (**b**) PCR confirmation of successful integration of the BirA-HA tag into the *PfMCMBP* locus in the PfMCMBP BirA-HA cell line. PCR was performed using the primers indicated in the schematic to confirm correct integration.

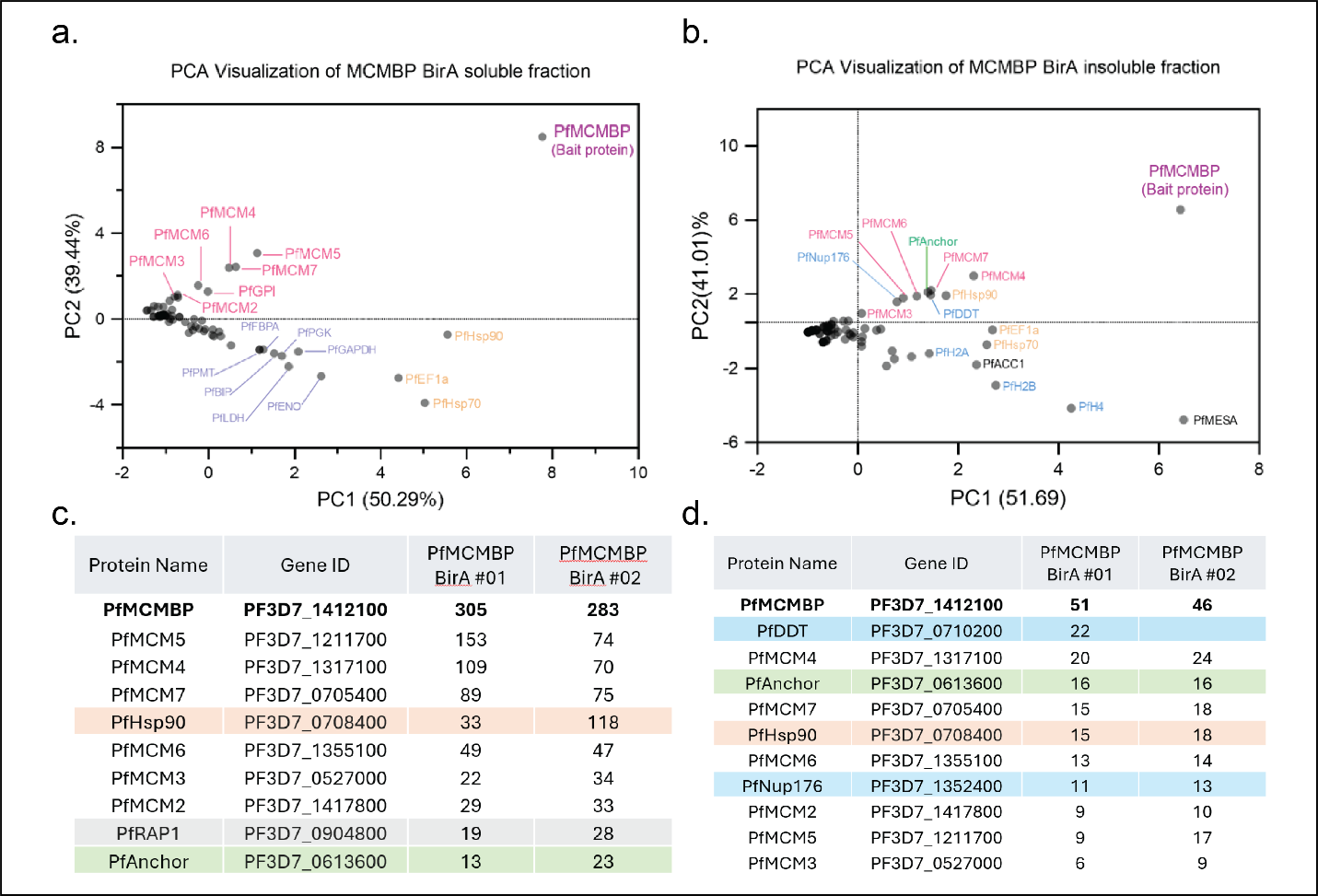

**Figure S2: Proximal labeling using PfMCMBP as bait identifies PfAnchor in both nuclear and cytosolic fractions.**

(**a-b**). Principal component analysis (PCA) of mass spectrometry data from proximal labeling using PfMCMBP as bait. (**a**) shows the nuclear (soluble) fraction, and (b) shows the cytosolic (insoluble) fraction. Labeled proteins represent enriched interactors identified in each fraction. (**c-d**). Tables listing the number of peptides detected for proteins uniquely present in the +biotin condition from the nuclear (**c**) and cytosolic (**d**) fractions in the PfMCMBP proximity labeling experiments.

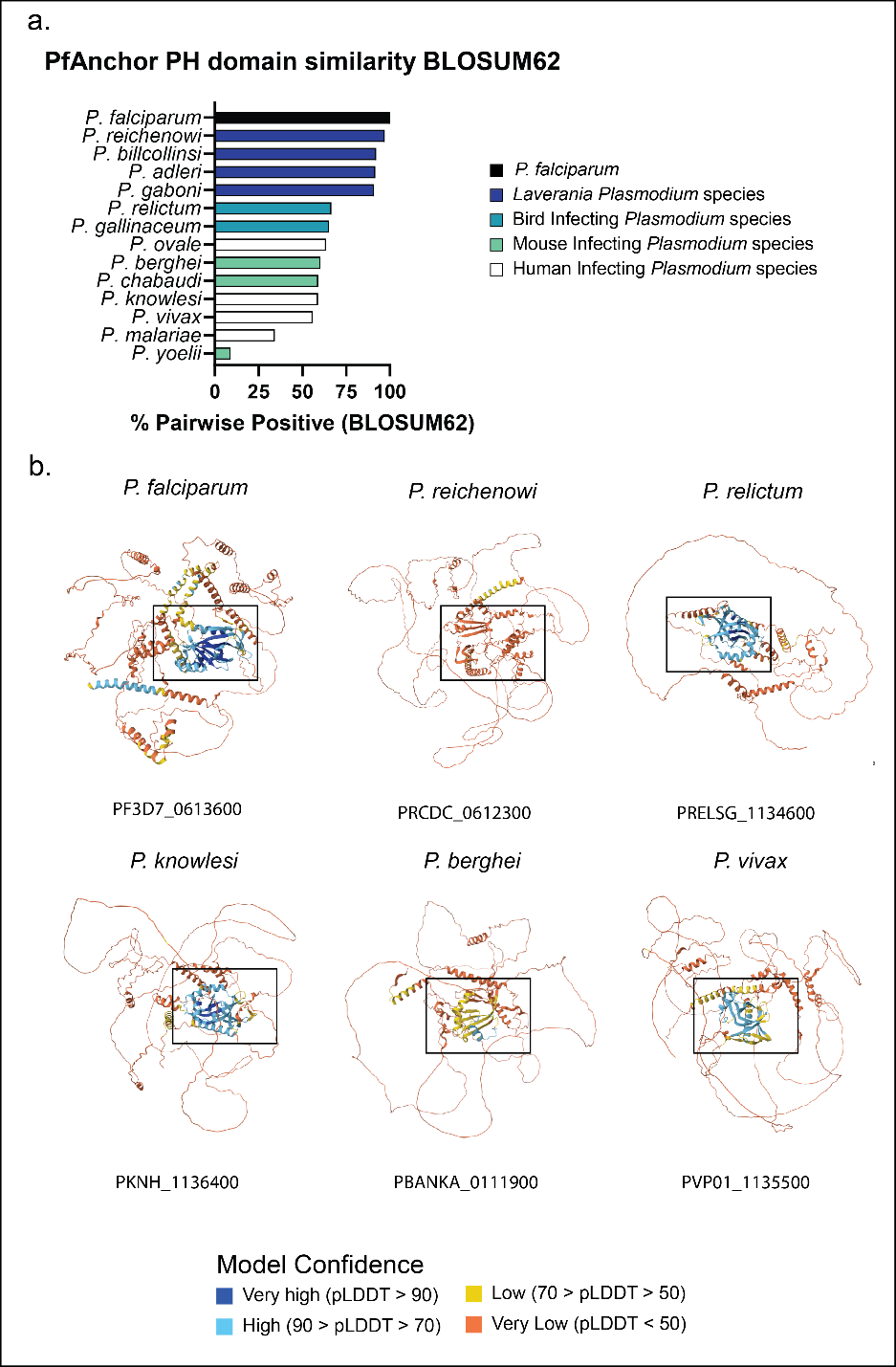

**Figure S3: Conservation of the PfAnchor PH domain across *Plasmodium* species.**
**a.** Pairwise sequence similarity of the predicted PH domain across *Plasmodium* species based on BLOSUM62 scoring. *Plasmodium* species from the *Laverania* subgenus exhibit the highest similarity to PfAnchor. **b.** Alphafold structure predictions of PfAnchor across various *Plasmodium* species. The predicted PH domain (black box) is structurally conserved and modeled with high confidence in most *Plasmodium* species.

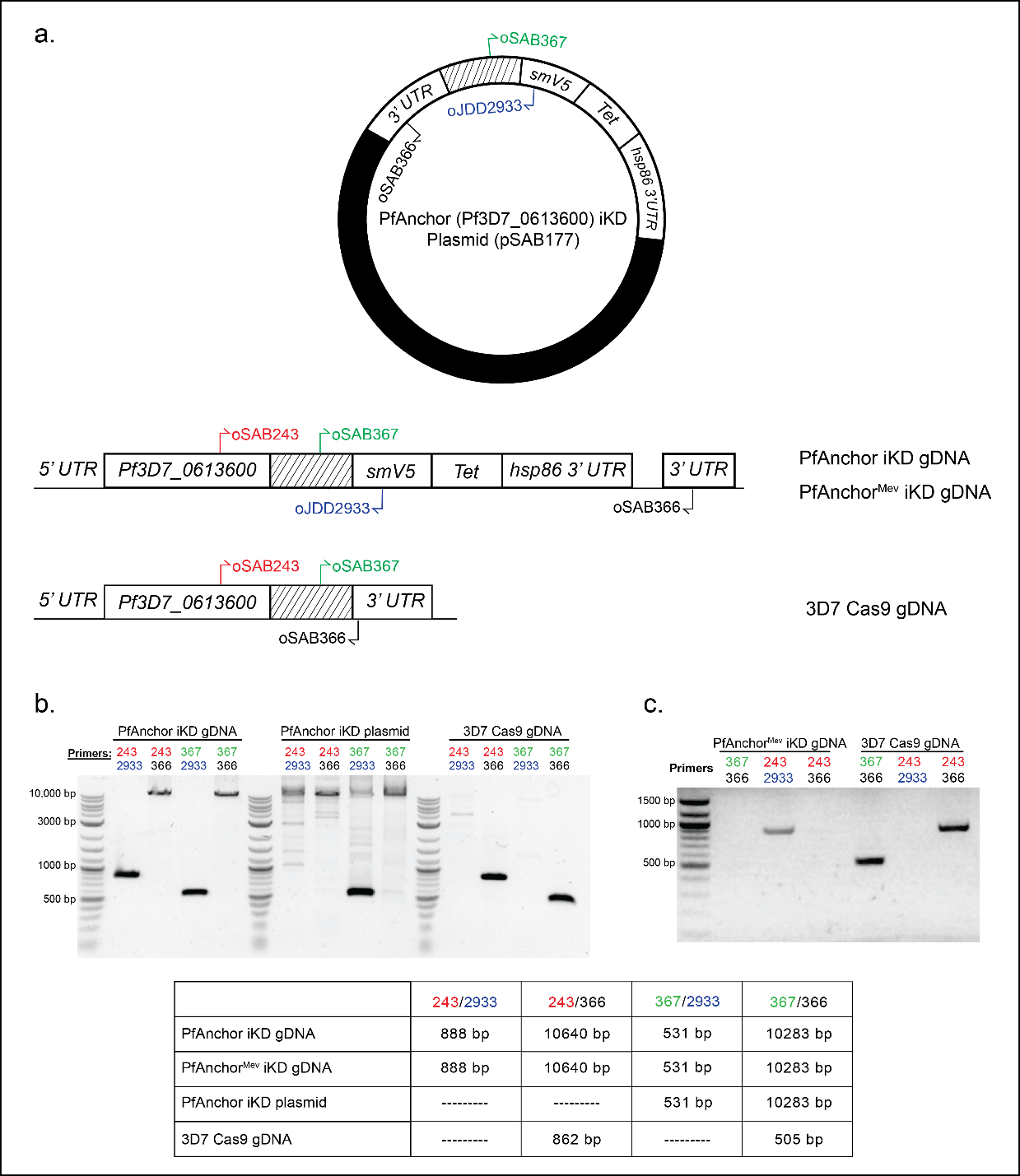

**Figure S4: Confirmation of PfAnchor iKD and PfAnchor^Mev^ iKD cell lines.**
**a.** Schematic representation of the PfAnchor (Pf3D7_0613600) locus before and after integration of the smV5-Tet tag. 3D7 Cas9 parasites were transfected with a plasmid containing a 3’ homology region of PfAnchor, followed by a C-terminal smV5-Tet tag. (**b-c**) PCR confirmation of successful integration of the smV5-Tet tag into the *PfAnchor* locus in both PfAnchor iKD and PfAnchor^Mev^ iKD cell lines using primers indicated in the schematic.

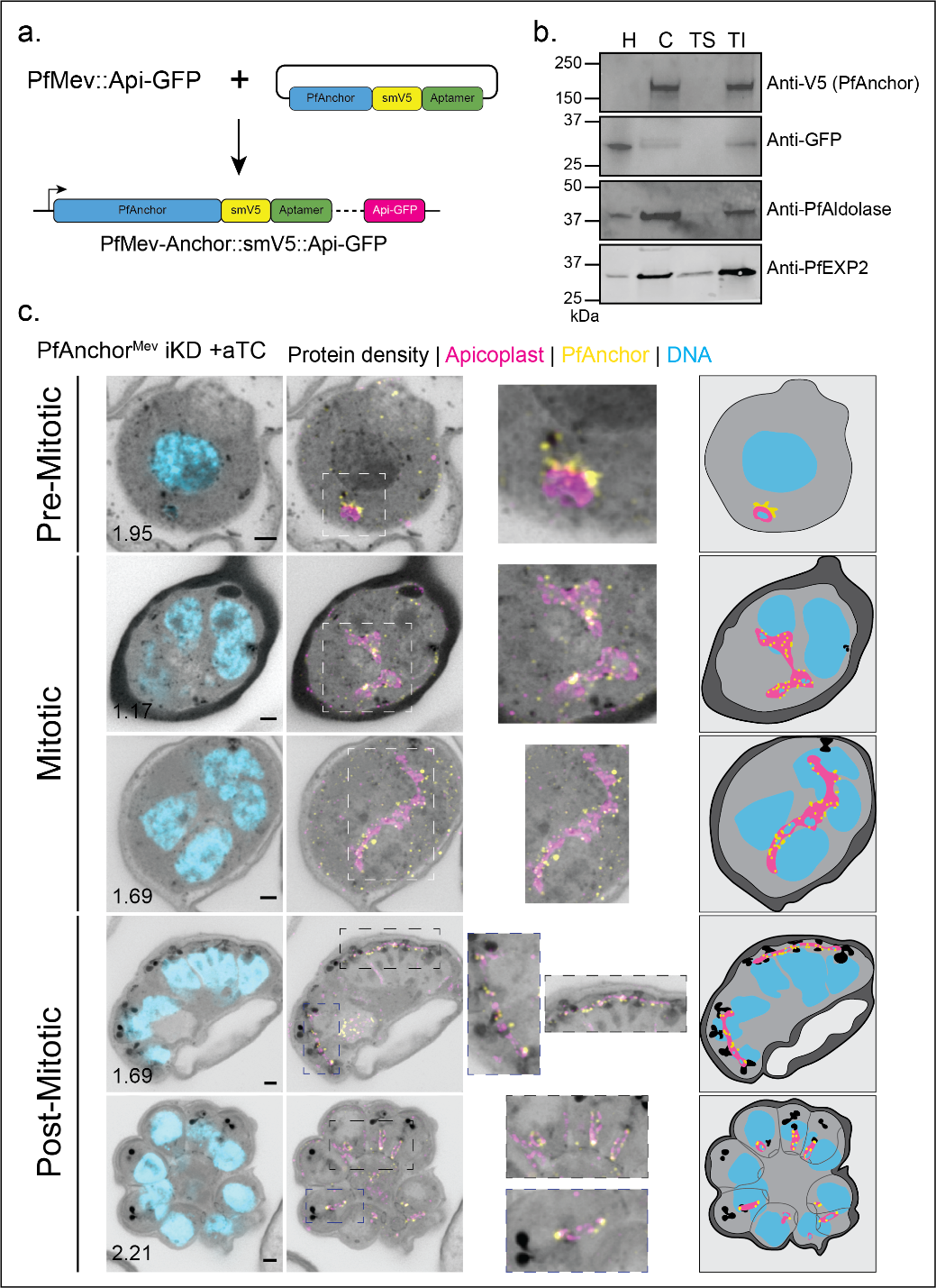

**Figure S5: PfAnchor is expressed throughout the asexual life cycle and localizes around the apicoplast.**

**a.** Schematic representation of the PfAnchor^Mev^ iKD cell line. This cell line expresses PfAnchor tagged with an smV5-Tet tag in the PfMev background, which also contains a GFP-tagged apicoplast marker **b.** Sodium carbonate fractionation assay demonstrating that PfAnchor is associated with a membrane. Fractions include hypotonic (unbound cytoplasmic proteins, H), carbonate (peripherally bound membrane proteins, C), Triton X-100 soluble (integral membrane proteins, TS), and Triton X-100 insoluble (GPI-anchored membrane proteins, TI). GFP (PfACP-GFP) was used as a non-membrane bound protein control, PfAldolase was used as a cytosolic and peripherally membrane bound control, and PfEXP2 was used as a integral membrane protein control. We note the lack of PfAnchor signal in the hypotonic (H) sample, indicating that PfAnchor is membrane associated. **c.** U-ExM images showing the localization of PfAnchor with respect to the apicoplast throughout the asexual blood-stage cycle. Scale bars: 2 µm, with image depth (µm) indicated.

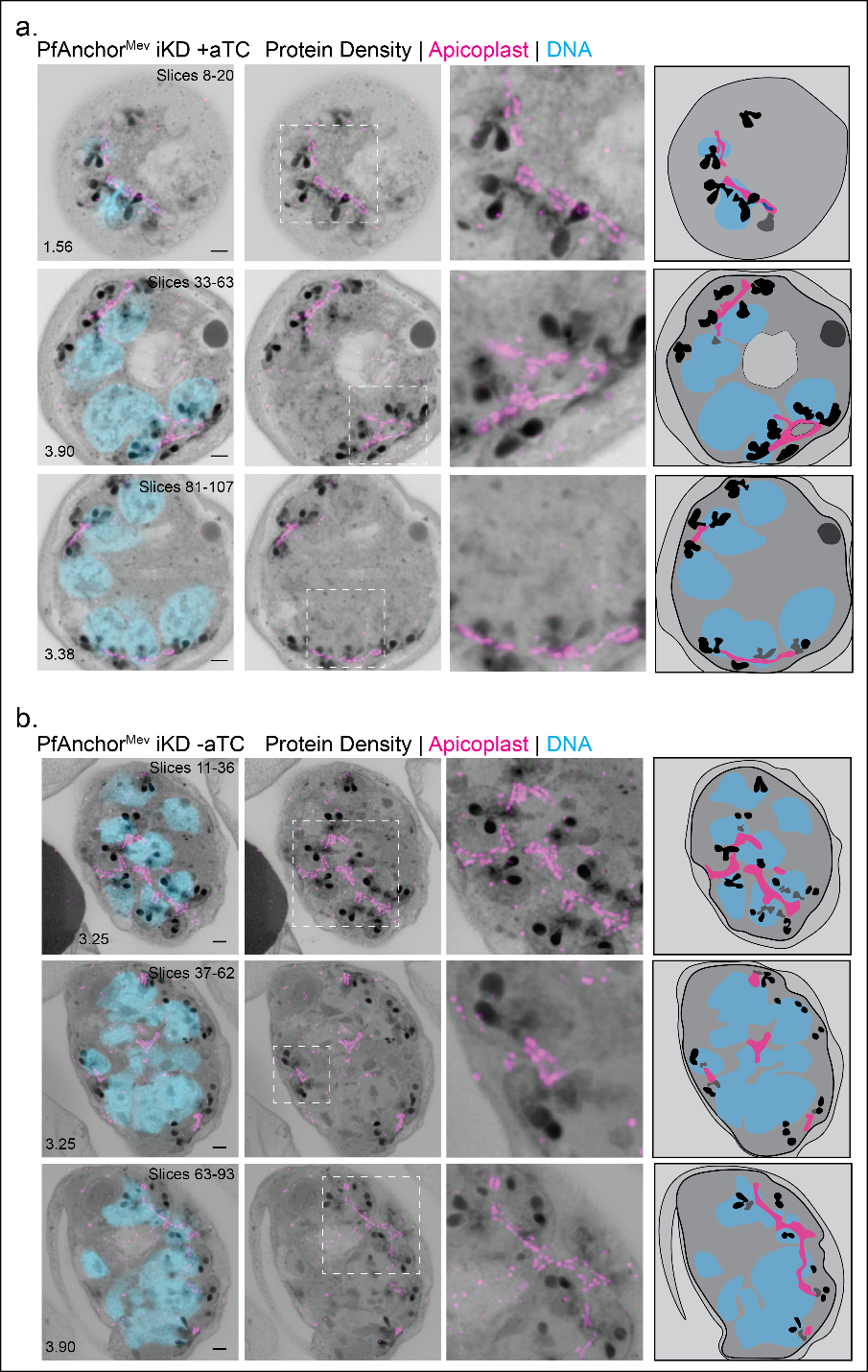

**Figure S6: PfAnchor knockdown does not affect apicoplast positioning or “crown” morphology upon fission.**

Representative U-ExM images showing the apicoplast’s characteristic “crown” morphology upon fission in PfAnchor-expressing (**+aTC, panel a**) and PfAnchor-deficient (**-aTC, panel b**) parasites. In both conditions, the apicoplast remains a single branching organelle that is associated with the centriolar plaques (CPs) during early segmentation, suggesting that PfAnchor depletion does not impair apicoplast positioning before cytokinesis. Apicoplasts (magenta), DNA (cyan), and protein density (grayscale) are shown. Scale bars: 2 µm.

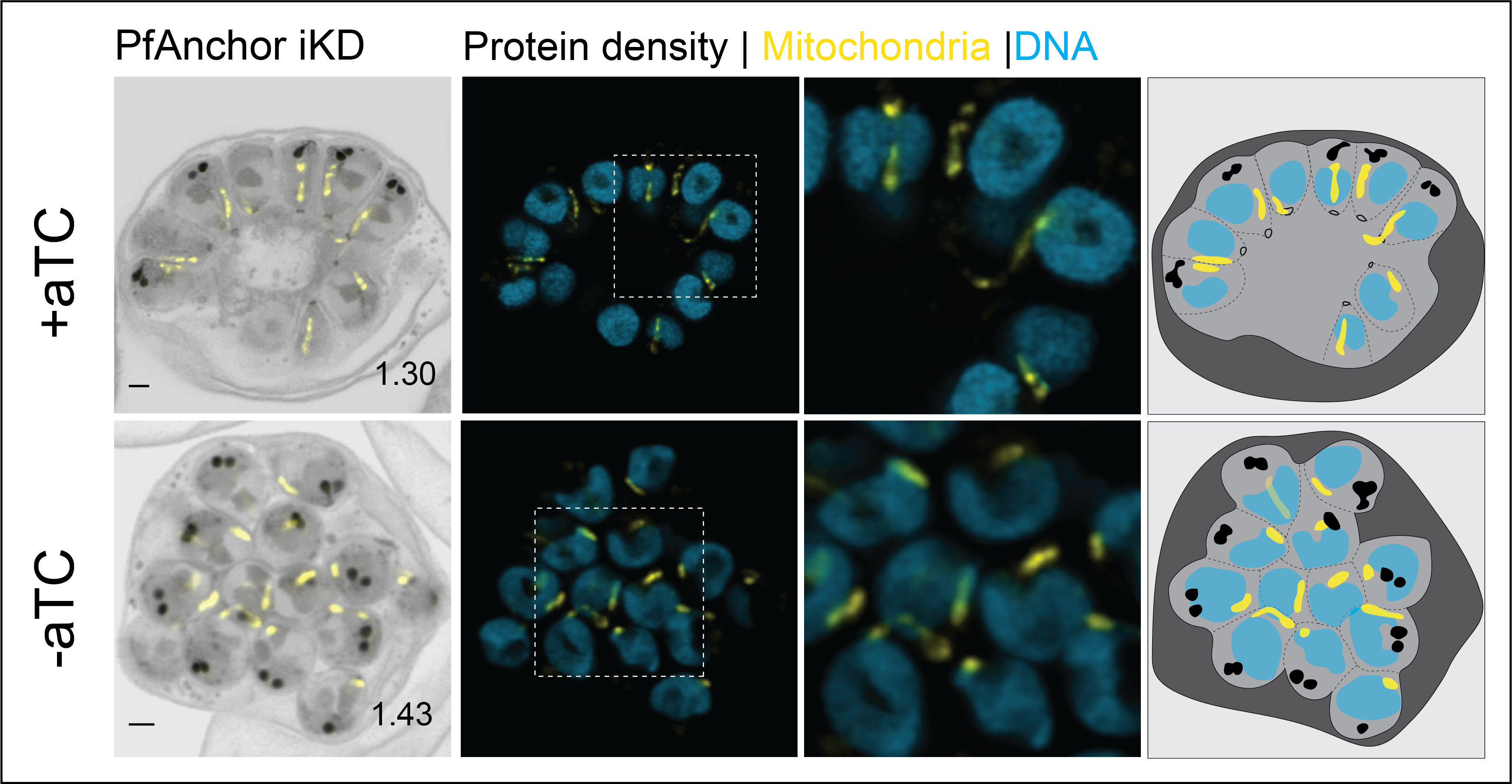

**Figure S7: PfAnchor knockdown does not affect mitochondrial division and inheritance.**

U-ExM images of parasites during cytokinesis, showing mitochondrial organization in PfAnchor-expressing (+aTC) and PfAnchor-deficient (–aTC) conditions. Mitochondrial signal (yellow) is observed to undergo proper segmentation in both conditions, indicating that PfAnchor depletion does not impair mitochondrial division and inheritance. Scale bars: 2 µm, with image depth (µm) indicated.

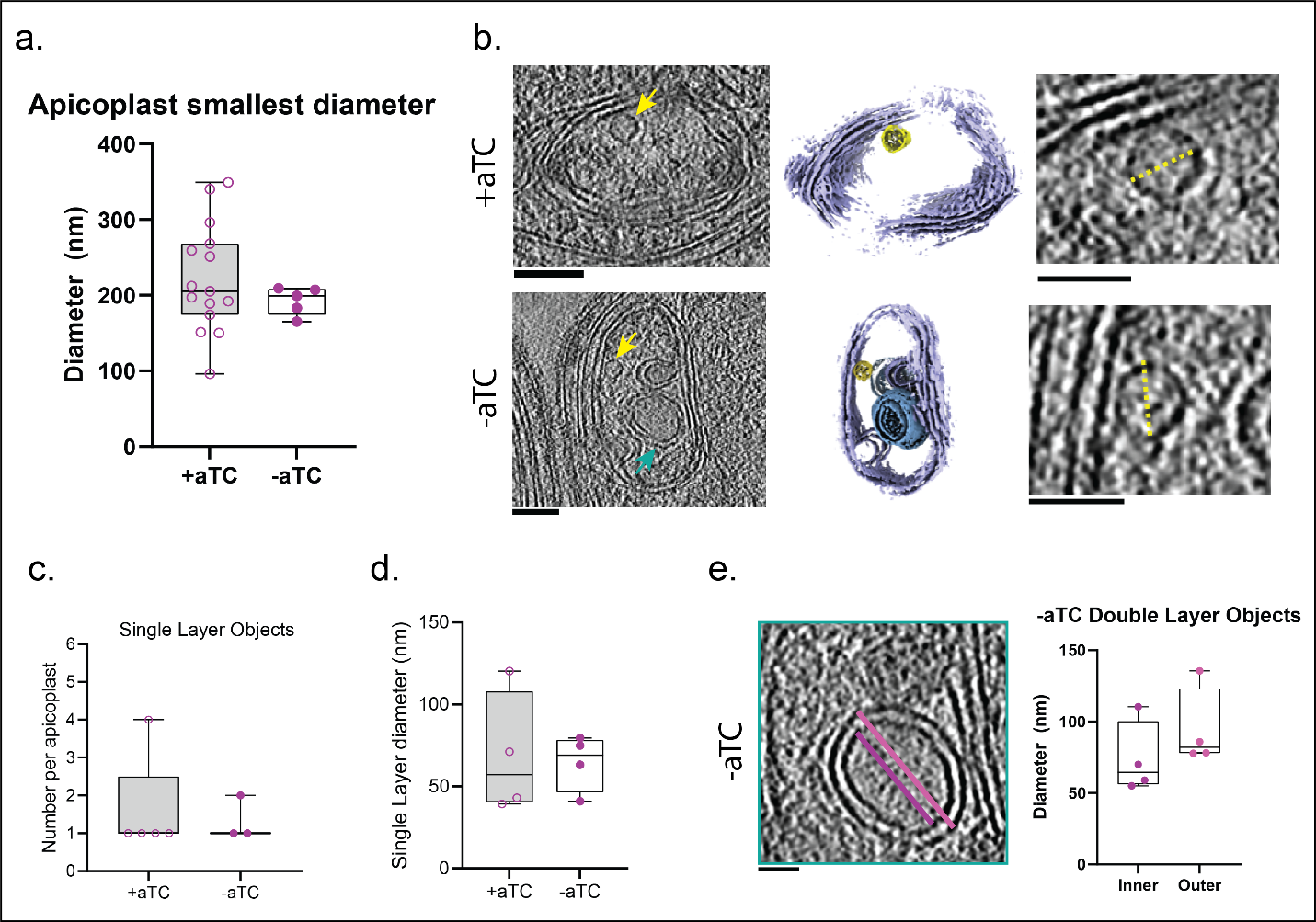

**Figure S8: Cryo-electron tomography (Cryo-ET) reveals internal apicoplast structures in PfAnchor-expressing and deficient parasites.**

**a.** Measurement of the smallest internal apicoplast diameter shows no significant difference between PfAnchor-expressing (+aTC) and PfAnchor-deficient (−aTC) parasites. **b.** Representative Cryo-ET images show single- and double-layered membranous structures inside the apicoplast lumen in PfAnchor-expressing and deficient parasites. Yellow arrows highlight these structures. **c.** Quantification of the number of single-layered membrane structures per apicoplast in PfAnchor-expressing and PfAnchor-deficient parasites. **d.** Quantification of the diameter of the single-layered membrane structures in the apicoplast lumen in PfAnchor-expressing and deficient parasites. **e.** Measurement of the inner (purple) and outer (magenta) diameters of double-layered membrane structures found inside the apicoplast lumen of PfAnchor-deficient parasites. All scale bars: 100 nm.

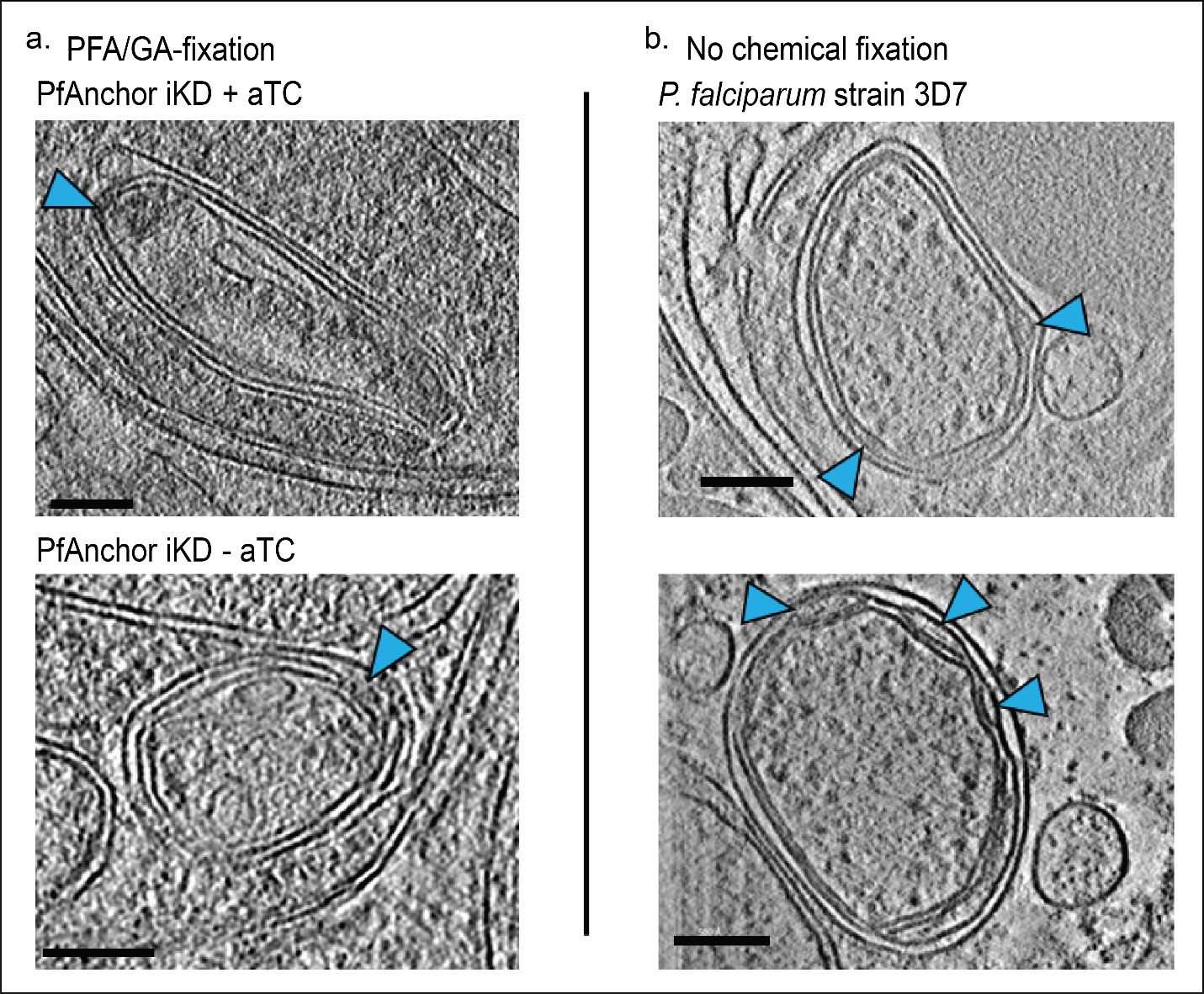

**Figure S9. Dense intermembrane buds are observed in both chemically fixed and unfixed cryo-preserved parasites.**

(**a**) Cryo-electron tomograms of PfAnchor iKD parasites prepared with paraformaldehyde/glutaraldehyde (PFA/GA) fixation prior to cryo-preservation (+aTC and –aTC conditions). (**b**) Cryo-electron tomograms of *P. falciparum* strain 3D7 prepared without chemical fixation prior to cryo-preservation (data from [1]). In both preparations, electron-dense structures bridging apicoplast membranes (blue arrowheads) are clearly observed, indicating that these “dense buds” are unlikely artifacts of chemical fixation. Scale bars: 100 nm.

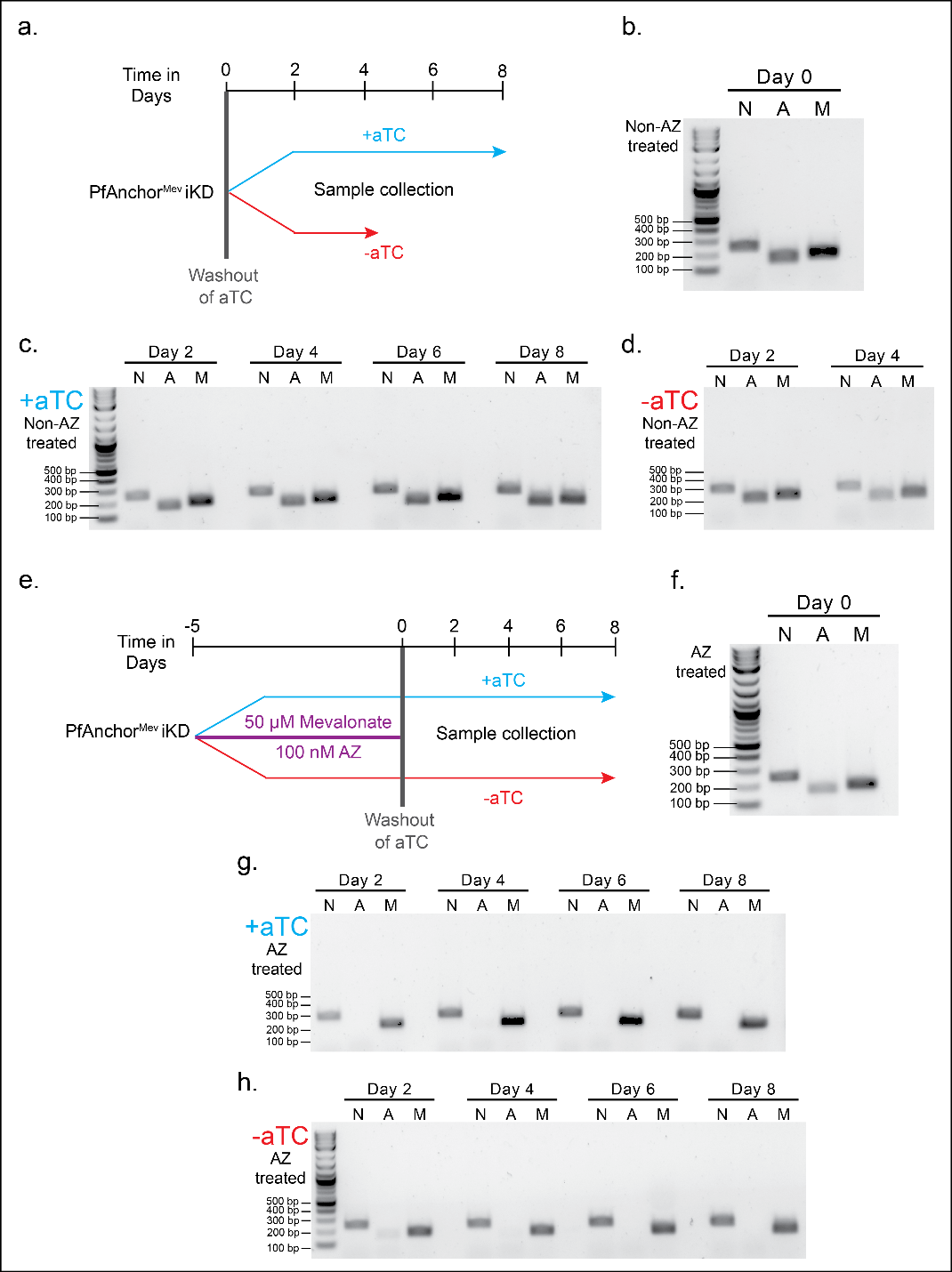

**Figure S10: Loss of the apicoplast genome upon azithromycin (AZ) treatment in PfAnchor^Mev^ iKD parasites**.

(**a, e**) Schematic representation of the experimental design. PfAnchorMev iKD parasites were grown in the presence or absence of aTC, with or without prior treatment with azithromycin (AZ) and mevalonate (Mev). For panels (**a–d**), parasites were cultured without mevalonate supplementation, thus mimicking PfAnchor depletion. After washout of aTC, samples were collected at the indicated time points. (**b, f**) DNA gel showing the presence of nuclear (N), apicoplast (A), and mitochondrial (M) genomes at Day 0. (**c, d**) Time-course DNA analysis of parasites without AZ treatment, in the absence (**c**) or presence (**d**) of aTC. In -aTC conditions, no parasites were detected after Day 4 due to the lack of PfAnchor and Mev, so no samples were collected beyond this point. (**g, h**) In AZ-treated parasites (with Mev), the apicoplast genome (A) was progressively lost over time in both +aTC and -aTC conditions, while nuclear (N) and mitochondrial (M) DNA remained detectable. Primers used: see Table S1

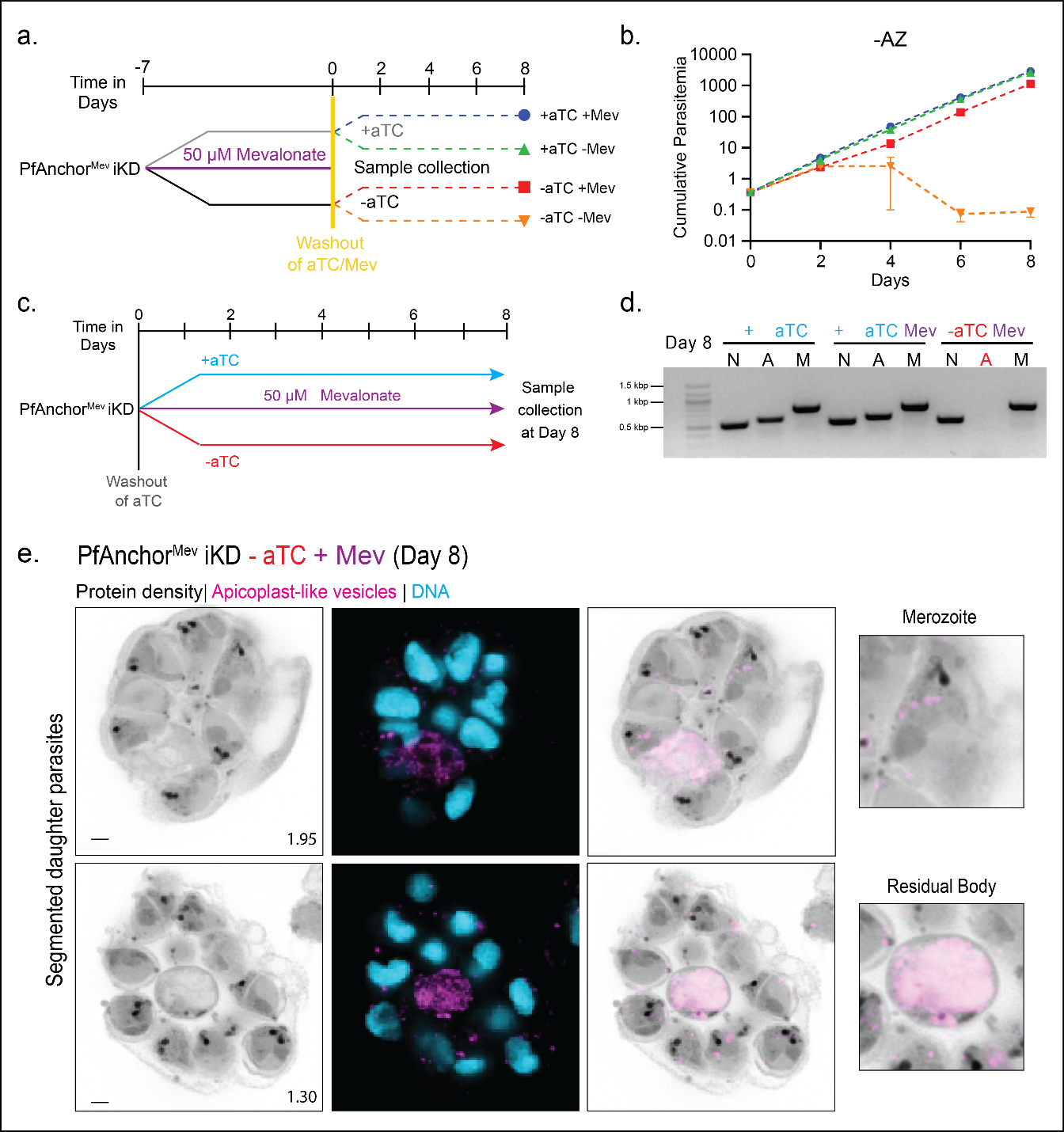

**Figure S11: Supplementation of PfAnchor-deficient PfAnchor^Mev^ iKD -AZ parasites with Mev rescues parasite growth after one cycle.**

**a.** Schematic of the experimental setup. PfAnchor^Mev^ iKD parasites were pre-treated with mevalonate (+Mev) for 7 days before prior to cell washing and maintaining under the indicated conditions (±aTC, ±Mev) for downstream analyses. **b.** Cumulative parasitemia of PfAnchor-expressing (+aTC) and PfAnchor-deficient (-aTC) parasites over 8 days in +AZ conditions. In AZ-treated parasites, disruption of the apicoplast resulted in the formation of apicoplast-like vesicles, which rescued the growth defect of PfAnchor-deficient parasites, but only in the presence of mevalonate (+Mev). Parasites lacking both PfAnchor and mevalonate (-Mev) failed to propagate. Parasites were sampled every 2 days for growth and PCR analysis, with cultures diluted 1:8 every 2 days to ensure continued replication. Data represent mean ± SD from 2 biological replicates in quadruplicate. **c.** Schematic of the experimental setup for confirming apicoplast negative parasites. PfAnchor sufficient and deficient parasites were supplemented with Mev for 8 days before taking samples for PCR and U-ExM analysis. **d.** PCR analysis of PfAnchor sufficient and deficient parasites at day 8. PfAnchor sufficient parasites with and without Mev supplementation showed robust signal of the apicoplast genome, while PfAnchor deficient parasites supplemented with Mev showed no apicoplast genome. **e.** U-ExM of PfAnchor-deficient parasites show the presence of apicoplast-like vesicles. These vesicles localize in the same manner as PfAnchor sufficient parasites treated with Azithromycin.

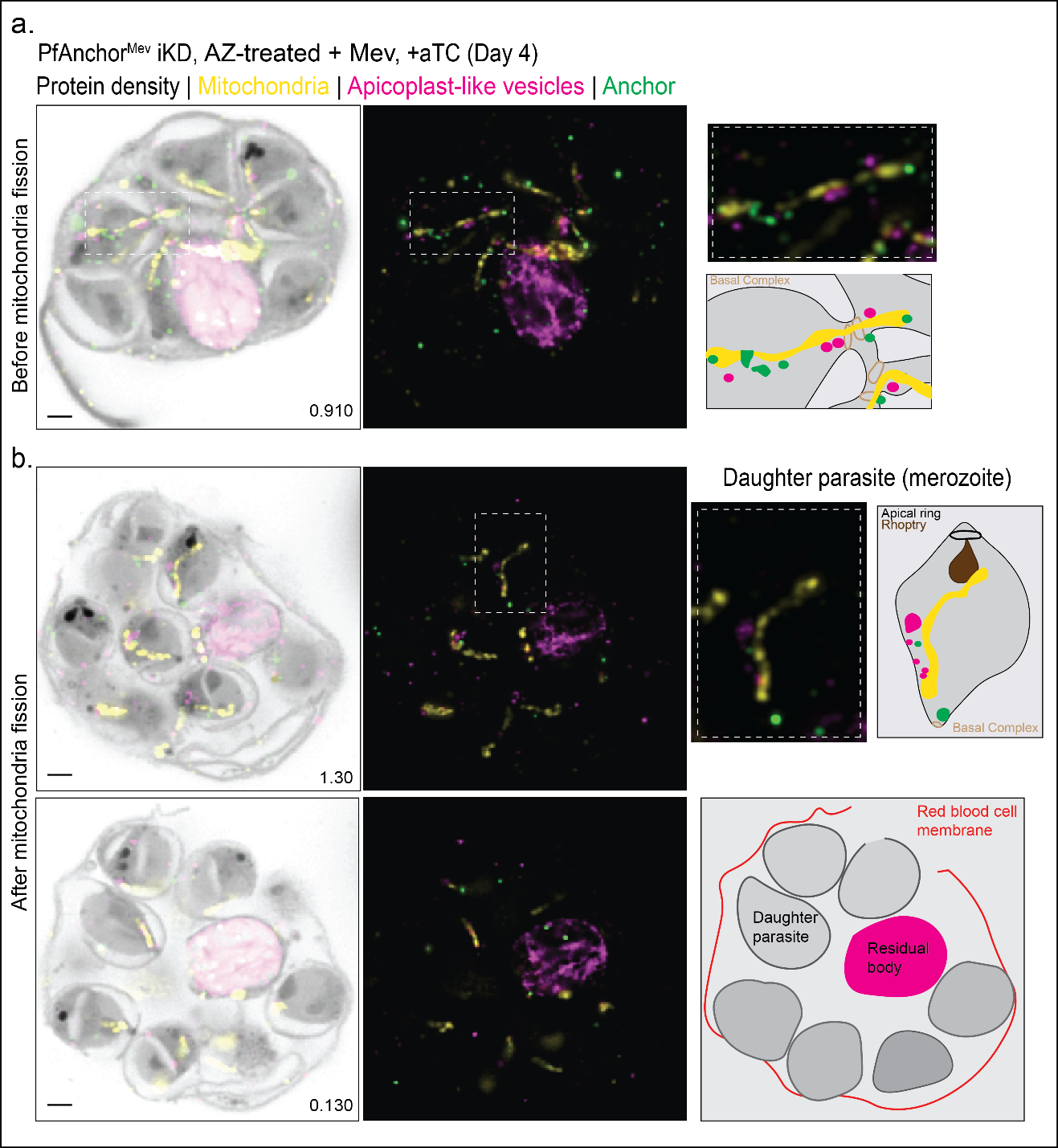

**Figure S12: Apicoplast-like vesicles retained within daughter parasites are predominantly associated with mitochondria.**

**a.** U-ExM image of a PfAnchor-expressing (+aTC) parasite treated with AZ/Mev, collected on Day 4 of the time-course experiment prior to mitochondrial fission. Most apicoplast-like vesicles (ACP-GFP, magenta) are localized to the residual body, with only a subset retained within developing daughter parasites. These retained vesicles are predominantly associated with mitochondria (yellow) and often colocalize with PfAnchor foci (green). Insets show zoomed-in views and a schematic representation of vesicle-mitochondria associations at the basal complex. **b**. U-ExM images of parasites collected after mitochondrial fission. Apicoplast-like vesicles are now more frequently retained within daughter parasites and remain associated with mitochondria. Right panels show higher magnification of the vesicle-mitochondrion association within an individual merozoite (top), and a schematic overview of parasite organization within the red blood cell (bottom). Protein density is shown in grayscale. Scale bars: 2 µm, with image depth (µm) indicated.

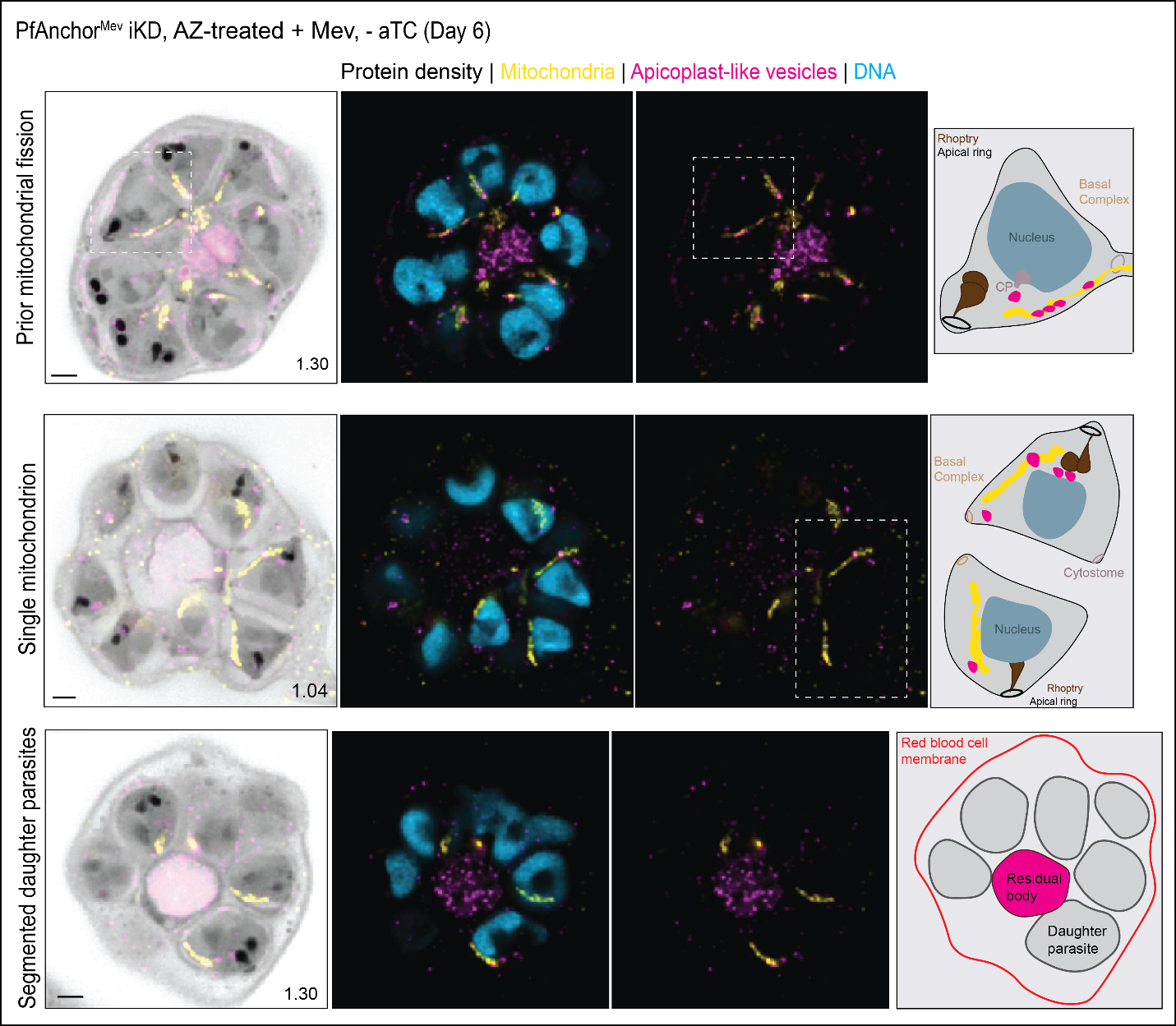

**Figure S13: PfAnchor is not required for apicoplast-like vesicle inheritance.**

U-ExM images of PfAnchor-deficient (-aTC) parasites treated with AZ/Mev, collected six days after aTC removal during cytokinesis. Similar to PfAnchor-expressing (+aTC) parasites, the majority of apicoplast-like vesicles (ACP-GFP, pink) localize to the residual body, while a subset remains associated with mitochondria (yellow) within daughter parasites. DNA is labeled in blue. Insets highlight vesicle-mitochondria interactions and their spatial organization within the developing parasite. These findings indicate that PfAnchor is not required for apicoplast-like vesicle inheritance. Scale bars: 2 µm, with image depth (µm) indicated.

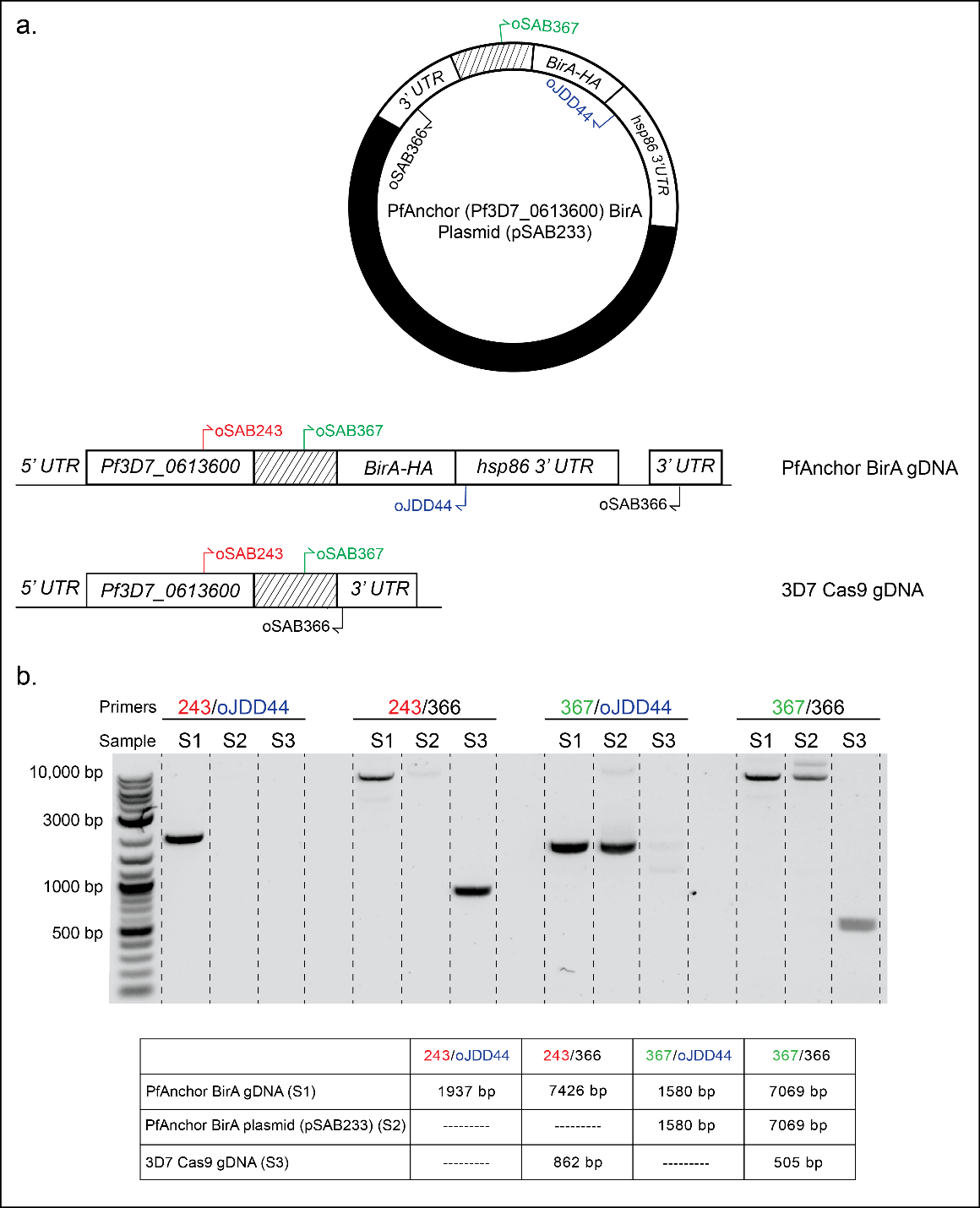

**Figure S14: Generation and confirmation of the PfAnchor BirA-HA cell line.**

**a.** Schematic representation of the PfAnchor (Pf3D7_0613600) locus before and after integration of the BirA-HA tag. 3D7 Cas9 parasites were transfected with pSAB233 containing a 3’ homology region (marked with hashed lines) of PfAnchor, followed by a C-terminal BirA-HA. (**b**) PCR confirmation of successful integration of the BirA-HA tag into the *PfAnchor* locus in the PfAnchor BirA-HA cell line. PCR was performed using the primers indicated in the schematic to confirm correct integration.

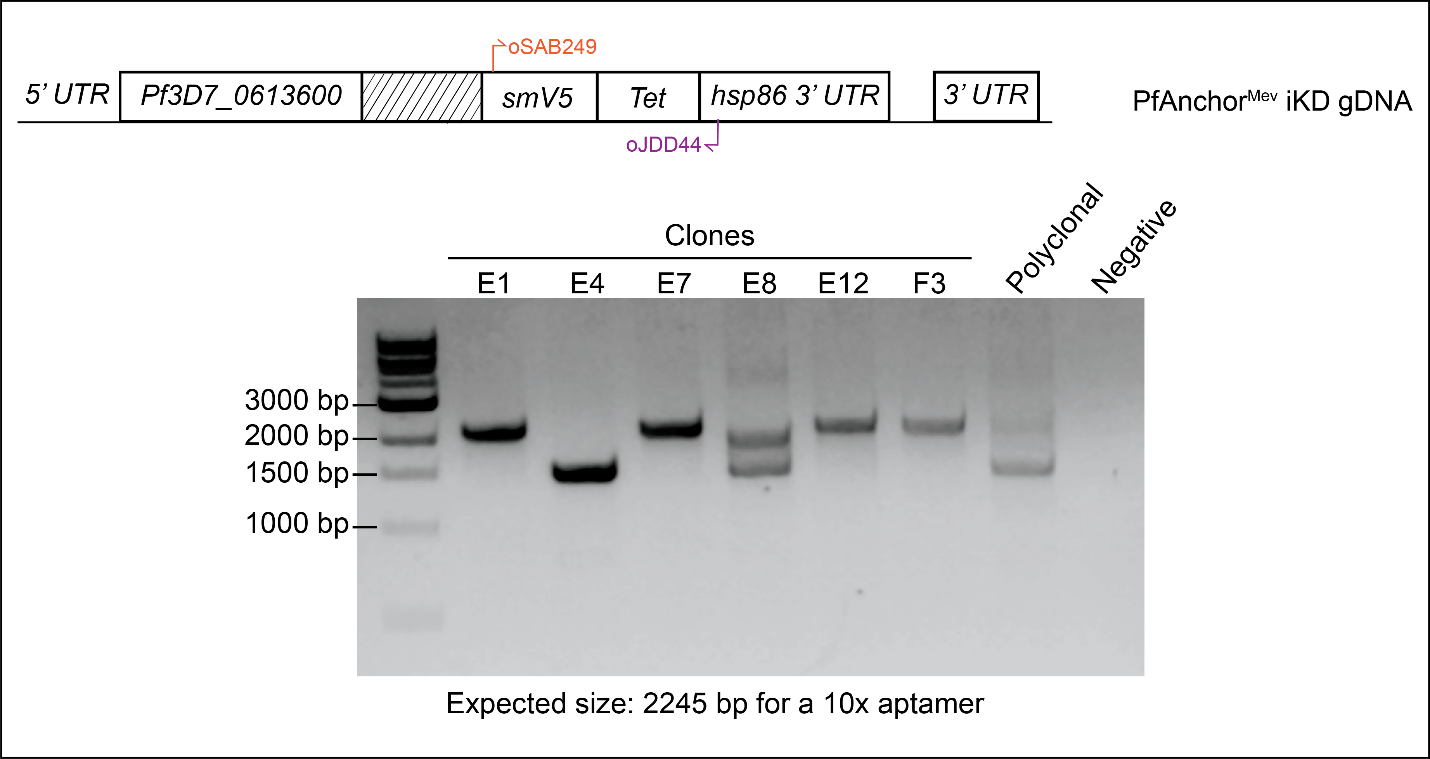

**Figure S15: PCR Confirmation of 10x aptamer in the PfAnchor^Mev^ iKD cell line.**

**a.** PfAnchor^Mev^ iKD parasites were subcloned by limiting dilution before parasite clones were selected and screened for a 10x aptamer for the TetR-DOZI aptamer knockdown system. PCR primers oSAB249 and oJDD44 were used to amplify genomic DNA. Clones E1, E7, E12, and F3 showed a 10x aptamer (2245 bp PCR product) and clone F3 was selected for future experiments.

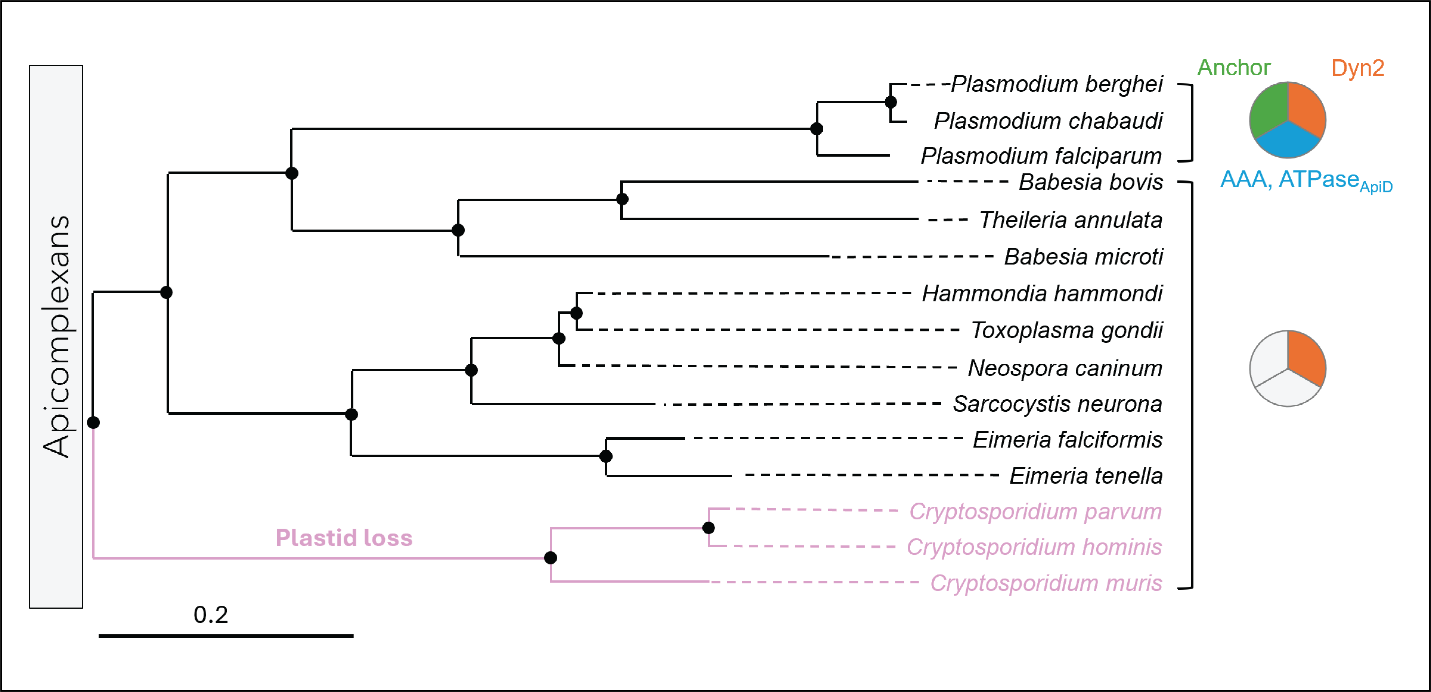

**Figure S16. Evolutionary distribution of candidate apicoplast divisome proteins across Apicomplexa.**
 We reproduced the relevant section of the published maximum-likelihood tree from Mathur et al., 2019 [2]. Topology and branch lengths follow the apicomplexan clade reported in that study, and black circles indicate well-supported nodes as defined by the authors. Onto this topology, we overlaid pie charts summarizing the presence (colored sectors) or absence (gray sectors) of candidate divisome proteins identified in Plasmodium falciparum, based on BLASTp searches against VEuPathDB reference genomes (see Methods for parameters). We also highlighted the Cryptosporidium lineage to reflect its secondary loss of plastids. Branch lengths are proportional to sequence divergence (scale bar = 0.2 substitutions/site).

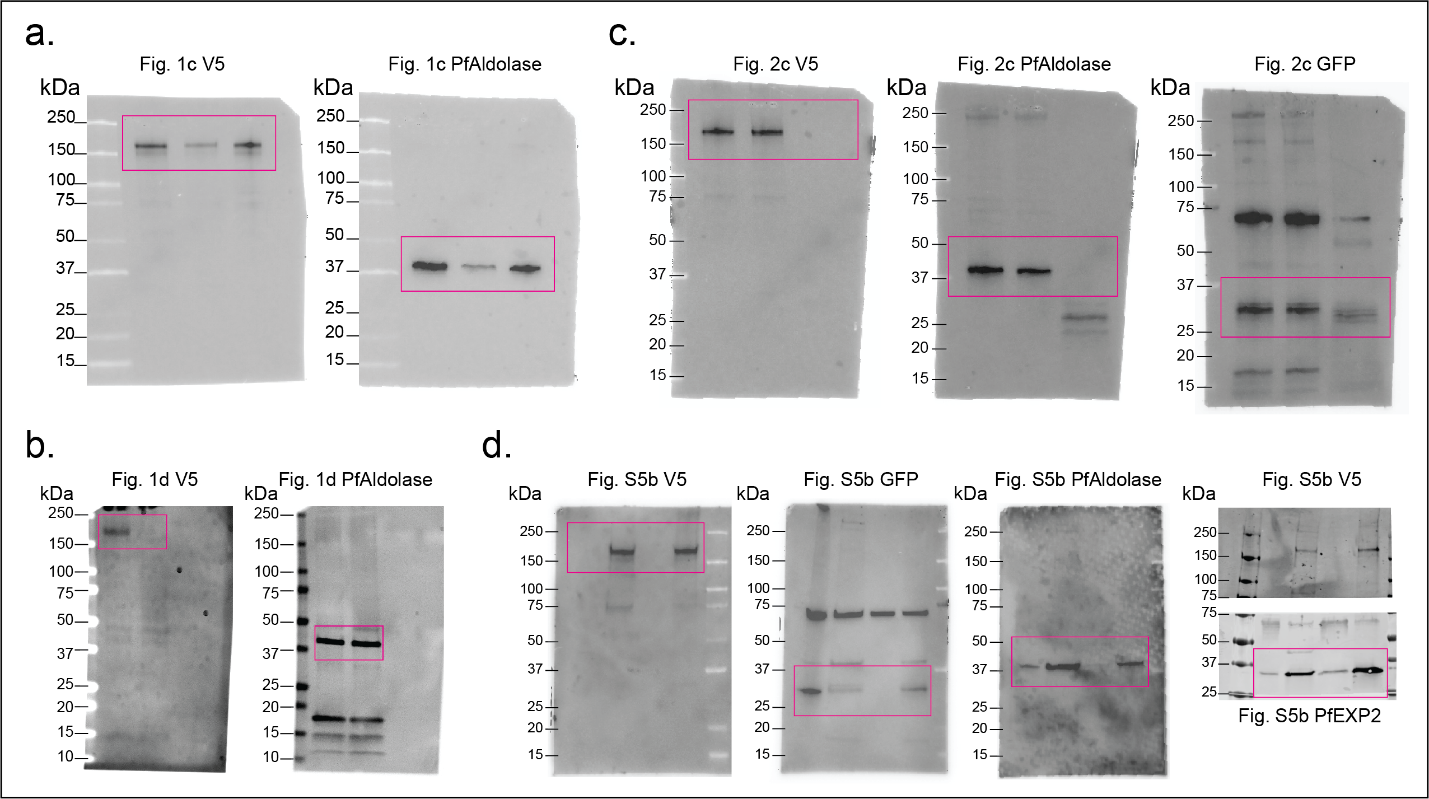

**Figure S17: Full-sized images of all immunoblots.** Full-sized immunoblots from Figure 1 (a, b), Figure 2 (c), and Figure S5 (d) are shown. The magenta box on each immunoblot indicates the cropped area displayed for the figures. The additional banding on the GFP stained blots is non-specific binding that we see in all of our GFP blots. For (d), we stained for V5 in multiple blots and saw similar staining profiles throughout our testing.

**Table S1: PCR primers used in this study.**

| **Primer** | **Use** | **Sequence** |
| --- | --- | --- |
| oJDD44 | PfAnchor aptamer hsp86 3' UTR reverse | TGGGGTGATGATAAAATGAAAG |
| oJDD56 | PfMCMBP reverse plasmid integration | ACACTTTATGCTTCCGGCTCGTATGTTGTG |
| oJDD2933 | PfAnchor integration smV5 reverse | CTGCTGCTGAGTACTATCAAGTC |
| oSAB243 | PfAnchor integration Anchor sequence forward | CAGATCAAAGAAGTGTCACAGGAATATC |
| oSAB249 | PfAnchor aptamer number smV5 forward | CCATGGATGGGAAAACCTATACCGAAC |
| oSAB356 | PfAnchor plasmid building 3HR forward | CATCGCGGCCGCAAGGGCACAAAATAAAATCAAGCATATATGGA |
| oSAB357 | PfAnchor plasmid building 5HR reverse | GATCCATGGCAGATGCTTGATGTTCCTCTCTTGTTTAAACACATGG |
| oSAB358 | PfAnchor guide plasmid building forward | ATATTGATGGATACACGTATTCAAGC |
| oSAB359 | PfAnchor guide plasmid building reverse | AAAACGCTTGAATACGTGTATCCATC |
| oSAB366 | PfAnchor integration 3' UTR reverse | CCATATATGCTTGATTTTATTTTGTGCCC |
| oSAB367 | PfAnchor integration plasmid forward | CATCAGCTTAATAATTTAGAGCTAGCTATAC |
| oSAB457 | PfSufB forward for Apicoplast genome analysis | CATGTAGCTATAGTAGAAATAATAGTAAAAGATTATGG |
| oSAB458 | PfSufB reverse for Apicoplast genome analysis | GACTCTGAAATACTTAAACCACGTTGC |
| oSAB480 | PfCox1 forward for Apicoplast genome analysis | CTTCATCTTTAAGAATAATTGCACAAGAAAATGTAAATC |
| oSAB481 | PfCox1 reverse for Apicoplast genome analysis | GGAAGCTTAGTATGGGTACATCATATGTAC |
| oSAB484 | PfGAPDH forward for Apicoplast genome analysis | ATCAAAGGGTGGTAAGGACTGG |
| oSAB485 | PfGAPDH reverse for Apicoplast genome analysis | AGTGGACCTTCAGCAGCTTTTT |
| oSAB486 | PfTufA forward for Apicoplast genome analysis | GATATTGATTCAGCTCCAGAAGAAA |
| oSAB487 | PfTufA reverse for Apicoplast genome analysis | ATATCCATTTGTGTGGCTCCTATAA |
| oSAB488 | PfCytb3 forward for Apicoplast genome analysis | AGATACATGCACGCAACAGG |
| oSAB489 | PfCytb3 reverse for Apicoplast genome analysis | TCATTTGACCCCATGGTAAGA |
| oSAB504 | PfMCMBP forward plasmid integration | TTCGGTAGCCTCATCAACGGACATATCT |
| oSAB505 | PfMCMBP forward plasmid integration | GACGTAATCGAAATTATTGGAATATATCGT |
| oSAB506 | PfMCMBP reverse plasmid integration | CATAATATAATCCCAGTGATCCCTA |
| LDH.F | PfLDH forward for Apicoplast genome analysis | GGAGATGTAGTTTTGTTCGATATTG |
| LDH.R | PfLDH reverse for Apicoplast genome analysis | CTTGTAAAGGGATACCACCTACAG |
| SufB.F | PfSufB forward for Apicoplast genome analysis | CATGTAGCTATAGTAGAAATAATAGTAAAAGA |
| SufB.R | PfSufB reverse for Apicoplast genome analysis | GACTCTGAAATACTTAAACCACGTTGC |
| Cox1.F | PfCox1 forward for Apicoplast genome analysis | CTTCATCTTTAAGAATAATTGCACAAGAAAATGTAAATC |
| Cox1.R | PfCox1 reverse for Apicoplast genome analysis | GGAAGCTTAGTATGGGTACATCATA |

**Table S2: Cell lines used in this study.**

| **Cell line** | **Description** |
| --- | --- |
| pSAB99 | 3D7 PfMCMBP^BioID^ |
| pSAB177 | 3D7 PfAnchor^smV5-Tet^ |
| PfMev-AnchorsmV5-Tet | NF54 PfMev-Anchor^smV5-Tet^ |
| pSAB233 | 3D7 PfAnchor^BioID^ |
| PfDyn2-3HAapt | Nf54 PfDyn2^3HA-Tet^ |

**Table S3: Primary antibodies, secondary antibodies, and stains used in this study.**

| **Antibody/Stain** | **Catalog number** | **Concentration** | **Use** |
| --- | --- | --- | --- |
| Rabbit Anti-GFP | OriGene (TP401) | 1:2000, 1:8000 | U-ExM, Western Blot |
| Chicken Anti-GFP | Abcam (ab13970) | 1:2000 | U-ExM |
| Anti-Aldolase | Abcam (ab207494) | 1:1000 | Western Blot |
| Anti-Histone H3 | Abcam (ab1791) | 1:2000 | Western Blot |
| Anti-V5 (Sy30-01) | biorad serotech MCA1360 | 1:250, 1:1000 | U-ExM, Western Blot |
| Anti-HSP60 | Invitrogen: PA5-34760 | 1:100 | U-ExM |
| Anti-EXP2 | Gift from Dr. Vasant Muralidarhan | 1:1000 | Western Blot |
| Anti-mouse IgG Alexa Fluor 555 | Thermo Fisher (A21428) | 1:500 | U-ExM |
| Anti-mouse IgG Alexa Fluor 568 | Thermo Fisher (A11004) | 1:500 | U-ExM |
| Anti-rabbit IgG Alexa Fluor 488 | Thermo Fisher (A11034) | 1:500 | U-ExM |
| Anti-rabbit IgG Alexa Fluor 555 | Thermo Fisher (A21428) | 1:500 | U-ExM |
| Anti-rabbit IgG Alexa Fluor 633 | Thermo Fisher (A21070) | 1:500 | U-ExM |
| Anti-rat IgG Alexa Fluor 488 | Thermo Fisher (A11006) | 1:500 | U-ExM |
| Anti-Chicken Alexa Fluor 488 | Thermo Fisher (A78948) | 1:500 | U-ExM |
| Anti-mouse Starbright Blue 520 | BioRad 12005867 | 1:2000 | Western Blot |
| Anti-Rabbit Starbright Blue 700 | BioRad 12004159 | 1:2000 | Western Blot |
| NHS ester Alexa Fluor 405 | Thermo Fisher (A30000) | 1:250 (8 µM) in DMSO | U-ExM |
| SYTOX Deep Red | Thermo Fisher (S11381) | 1:1000 (1 µM) in DMSO | U-ExM |
| Anti-Mouse HRP | Thermo Fisher (31430) | 1:8000 | Western Blot |
| Anti-Rabbit HRP | Thermo Fisher (31460) | 1:8000 | Western Blot |
| Anti-PfAldolase | Abcam (AB38905) | 1:8000 | Western Blot |
